# ZNF143 binds DNA and stimulates transcription initiation to activate and repress direct target genes

**DOI:** 10.1101/2024.05.13.594008

**Authors:** Jinhong Dong, Thomas G. Scott, Rudradeep Mukherjee, Michael J. Guertin

## Abstract

Transcription factors bind to sequence motifs and act as activators or repressors. Transcription factors interface with a constellation of accessory cofactors to regulate distinct mechanistic steps to regulate transcription. We rapidly degraded the essential and ubiquitously expressed transcription factor ZNF143 to determine its function in the transcription cycle. ZNF143 facilitates RNA Polymerase initiation and activates gene expression. ZNF143 binds the promoter of nearly all its activated target genes. ZNF143 also binds near the site of genic transcription initiation to directly repress a subset of genes. Although ZNF143 stimulates initiation at ZNF143-repressed genes (i.e. those that increase expression upon ZNF143 depletion), the molecular context of binding leads to *cis* repression. ZNF143 competes with other more efficient activators for promoter access, physically occludes transcription initiation sites and promoter-proximal sequence elements, and acts as a molecular roadblock to RNA Polymerases during early elongation. The term *context specific* is often invoked to describe transcription factors that have both activation and repression functions. We define the context and molecular mechanisms of ZNF143-mediated *cis* activation and repression.

## Introduction

Transcription factors bind directly to DNA and recruit cofactors and RNA Polymerase to regulate gene expression throughout the lifetime of all organisms. Transcription factors specify tissue patterning throughout development and their function is necessary to maintain cellular and organismal homeostasis (Lewis 1978; Nüsslein-Volhard and Wieschaus 1980). Sequence-specific transcription factors are comprised of DNA binding domains that recognize degenerate sequence motifs and effector domains that interface with cofactors that specialize in regulatory roles such as chromatin remodeling, initiation, pause release, and elongation. Just as cofactors are specialists, transcription factors can specialize by predominantly regulating specific steps in transcription (Duarte et al. 2016; Scholes et al. 2017; Scott et al. 2024). Factors achieve this specificity by preferentially recruiting certain cofactors through their effector domains. We propose that it is critical to classify transcription factors by their molecular function, as opposed to broad activator and repressor classes, in order to understand the context specificity of gene regulation. For example, recruiting a transcription factor that specializes in RNA Polymerase pause release to a gene may have little effect on transcription if redundant pause-release factors are already present; at this gene a factor that recruits components of the pre-initiation complex may cause potent activation. This work provides a framework for systematically determining the molecular functions of transcription factors by stimulating their rapid depletion and quantifying changes in RNA polymerase distribution at direct target genes in the minutes following transcription factor depletion.

Despite advances in understanding the mechanisms of transcription and developments in the systems biology of gene regulation field, accurately predicting direct target genes of transcription factors is a challenge. This struggle to predict target genes is in part because of the lack of reliable input data into predictive models. Researchers cannot experimentally identify a comprehensive set of primary response genes for the vast majority of transcription factors because they cannot rapidly induce or rapidly inhibit their activity. We know the most about transcription factors that are rapidly activated by acute environmental changes such as heat stress or nuclear receptor factors that are induced by addition of steroids (Duarte et al. 2016; Guertin et al. 2010; Kumar and Chambon 1988; Kumar et al. 1987; Westwood et al. 1991). Many models rely on input data where a transcription factor is depleted chronically for days or the lifetime of a cell, so it is impossible to discriminate target genes from the complex cascade of transcription dysregulation that follows. However, recent development of rapidly inducible degron systems democratizes the study of transcription factors that are not easily activated or inhibited (Nabet et al. 2020, 2018; Nishimura et al. 2009). Here, we C-terminally tagged all endogenous copies of the ubiquitous and essential transcription factor ZNF143 with FKBP^F36V^ in HEK-293T cells. We rapidly depleted ZNF143 by adding dTAG^V^-1 to study its molecular function and define ZNF143’s target genes.

The Xenopus homolog of ZNF143 was cloned and described three decades ago (Schuster et al. 1995). This original report characterized the binding site and activator function of ZNF143 (contemporaneously termed Staf) using reporter assays (Schuster et al. 1995). Subsequent transcriptional profiling upon chronic ZNF143 depletion identified many up regulated and down regulated genes (Ngondo-Mbongo et al. 2013); more genes were up regulated and the authors concluded that ZNF143 is primarily an activator while noting that ZNF143 may be involved in repression. Our results confirm that ZNF143 binds DNA to activate local genes and defines the mechanism of ZNF143 mediated activation. Furthermore, we define several molecular mechanisms through which a canonical activator can paradoxically retain its activator function while directly repressing target genes in *cis*. Although *cis* repression may only account for up to 30% of direct ZNF143-repressed targets, alternative mechanisms of immediate indirect repression, such as relieving competition for cofactors (Guertin et al. 2014; Schmidt et al. 2016, 2015), may explain why additional genes are repressed indirectly.

Many publications erroneously report that ZNF143 has a prominent role in chromatin looping (Bailey et al. 2015a; Heidari et al. 2014; Liu et al. 2022; Yang et al. 2017; Zhang et al. 2016, 2024). However, two recent studies found that ZNF143 has no role in chromatin looping (Magnitov et al. 2024; Narducci and Hansen 2024). These groups convincingly determine that the reason ZNF143 was ascribed a looping role is because a commonly used ZNF143 antibody crossreacts with the looping transcription factor CTCF (Magnitov et al. 2024; Narducci and Hansen 2024). ZNF143’s importance is highlighted by the rarity of characterized human disease with ZNF143 mutations because a lack of ZNF143 function is potentially incompatible with life. A case report of a child with missense and premature stop codon ZNF143 alleles exhibited deficiencies related to the lack of viable vitamin B12 cofactors and presented with global developmental delay (Pupavac et al. 2016). The authors linked the dysregulation of the ZNF143-regulated mitochondrial protein Methylmalonyl-CoA mutase to the observed signs and symptoms of vitamin B12-deficiency (Pupavac et al. 2016). This is consistent with the role of ZNF143 regulating nuclear encoded mitochondrial genes (Magnitov et al. 2024; Narducci and Hansen 2024). The patient also exhibited a range of symptoms that were not directly explainable by the effect of this mutation on vitamin B12 metabolism. Recent work shows that ZNF143 depletion in mouse embryo models limits the ability of stem cells to differentiate into their final lineages (Magnitov et al. 2024), which demonstrates ZNF143’s important role in early development and is a plausible explanation for the constellation of symptoms experienced by this patient. The only other reported case of ZNF143 in human disease is a family with a history of inherited endothelial corneal dystrophy, which results in premature thickening of the cornea and requires surgery at a young age to preserve vision (Kim et al. 2019). This family’s ZNF143 missense mutation is distinct from those of the previous case and appears to be only associated with the corneal symptoms, although vitamin metabolism abnormalities or symptoms affecting other organ systems were not reported. The varied manifestations and rarity of ZNF143 mutations in human populations make it difficult to study, but also demonstrates its importance to a wide variety of pathways that contribute to health and cell viability. For an essential protein such as ZNF143, rapidly inducible perturbation systems are generally more powerful when studying molecular mechanisms because molecular phenotypes can be quantified in the minutes and hours after losing the proteins function before downstream effects or compensatory mechanisms come into play.

## Results

### ZNF143 is rapidly degraded from DNA within 30 minutes

To determine the molecular function of ZNF143, we engineered HEK-293T cells to express all alleles of ZNF143 tagged with a C-terminal inducible dTAG degron system, facilitating rapid and complete protein degradation (Nabet et al. 2018). Endogenous tagging of ZNF143 causes a modest decrease in protein levels (Figure 1A lanes 1&2). Quantitative western blots indicate that less that 10% of ZNF143 remains after 15 minutes of dTAG^V^-1 treatment and ZNF143 is not detected at 30-90 minutes after induced degradation. We profiled ZNF143 binding genome-wide by ChIP-seq before and after 30 minutes of degradation to quantify depletion on chromatin. On the same scale, ZNF143 is not detectable after 30 minutes of degradation, but digital over-exposure of the heatmaps indicates that less than 5% of ZNF143 remains bound to DNA (Figure S1A). This cell line represents a powerful resource for investigating the molecular functions of ZNF143 and directly identifying ZNF143 target genes.

**Fig. 1.**
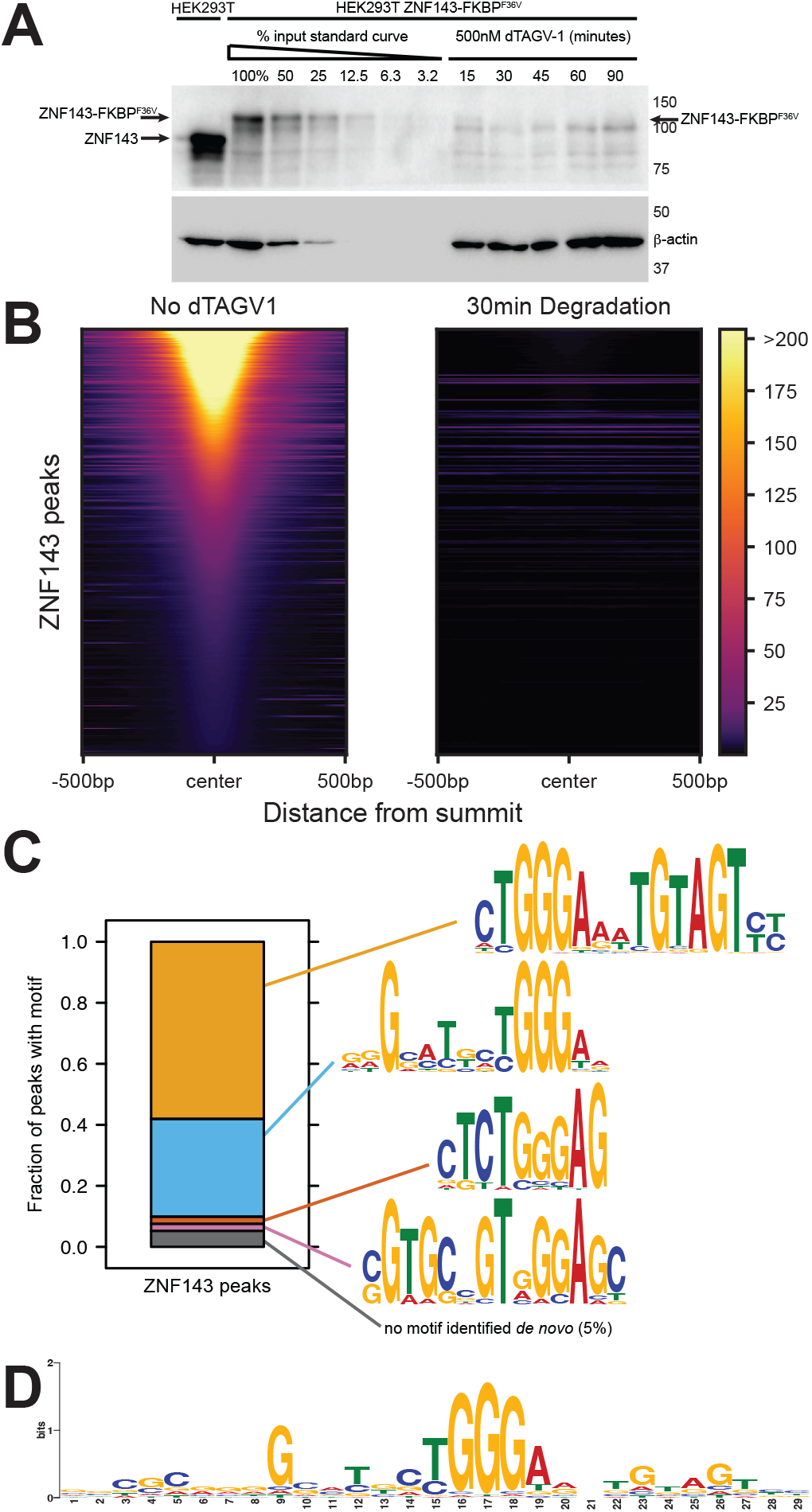
ZNF143 is degraded and off chromatin within 30 minutes of dTAG^V^-1-induced degradation. A) Quantitative western blots indicate that FKBPdegron^F36V^-tagged ZNF143 (lane 2) is expressed at comparable levels to untagged ZNF143 (lane 1). Lanes 2-7 are dilutions of the untreated and tagged cells line. Lanes 8-12 are a time series of degradation. Note that the arrow represents ZNF143 and there is a nonspecific band that is unchanged directly below the arrow. B) A heatmap of all ZNF143 ChIP-seq peaks indicates that ZNF143 is off chromatin after 30 minutes of degradation. C) Iterative *de novo* motif analysis identified four ZNF143 motifs that are anchored on a central TGGGA sequence. Ninety-five percent of ZNF143 peaks have at least one motif that conforms to a *de novo*-identified motif with a p-value of 0.0005 of lower. D) The 29 base seqLogo represents the average ZNF143 binding site, where substantial degeneracy is tolerated outside the core TGGGA postion.

### ZNF143 binds a degenerate 29 base motif at each binding site in the genome

Next, we conducted iterative and exhaustive motif analysis of ZNF143 ChIP-seq peaks to systematically identify and characterize the diverse sequence motifs that ZNF143 binds in the genome. We performed *de novo* motif analysis within the 4682 ZNF143 ChIP-seq peaks and identified a canonical ZNF143 sequence motif within 58% peaks (Figure 1C). ZNF143’s seven zinc fingers are known to bind a wider region (Zhang et al. 2024), so we performed the same analysis on remaining peaks without the first identified motif. We found a second ZNF143 motif variant and two more iterations of this process identified other ZNF143 motifs (Figure 1C). Ninety-five percent of the peaks had a *de novo*-identified motif that was clearly a ZNF143 motif variant (Figure 1C). These findings demonstrate ZNF143’s flexibility in binding; it can accommodate a wide range of sequences beyond the core **TGGGA** sequence that is recognized by zinc fingers 5 and 6 (Zhang et al. 2024). By anchoring the analysis on this core sequence, we calculated the average frequency of each nucleotide (A, C, G, T) within a 100-base range, highlighting how ZNF143’s binding preferences extend into the surrounding genomic landscape. The nucleotide frequencies stabilize to the genomic background outside a 29-mer core ZNF143 motif. This 29mer core is similar to the biochemically determined 27-mer ZNF143 core motif found previously (Zhang et al. 2024). We generated a composite motif using the frequencies in this window (Figure 1D). We did not identify a motif *de novo* in 5% of binding sites; however, these peaks remain sensitive to ZNF143 depletion and retain potential functionality. Low-affinity binding sites of transcription factors, which do not strictly conform to a consensus motif, are increasingly recognized as having a critical role in gene regulation and development (Jindal et al. 2023; Lim et al. 2024). We determined the precise ZNF143 binding positions within each ChIP-seq peak by assigning the motif with the lowest p-value in the peak as the inferred 29 base binding site (Figure S1B). All subsequent references and analyses concerning ZNF143 binding focus on the 29-mer ZNF143 recognition sequences within the ChIP-seq peaks. These analyses confirm the DNA-binding function of ZNF143 and indicate that ZNF143 binds a sequence-degenerate 29 base wide motif within chromatin.

### ZNF143 maintains chromatin accessibility at a minority of binding sites

To investigate the impact of ZNF143 on chromatin accessibility, we conducted ATAC-seq analysis before and after 30 minutes of ZNF143 depletion. Fewer than 500 ATAC-seq peaks significantly (FDR < 0.1) change accessibility after 30 minutes of ZNF143 depletion (Figure 2A). Over 99% of the significantly changed ATAC peaks decrease accessibility and 94% of the decreased accessibility peaks overlap ZNF143 binding sites (Figure 2B&C). Although the vast majority of decreased ATAC peaks are directly regulated by ZNF143 binding, the inverse is not true. Only a minority (10%; 479/4684) of ZNF143 peaks significantly reduce chromatin accessibility upon ZNF143 ablation from chromatin. These results underscore that while ZNF143 maintains open chromatin at certain loci, its influence is not uniform across all binding sites, suggesting that regulating chromatin structure is not its primary function.

**Fig. 2.**
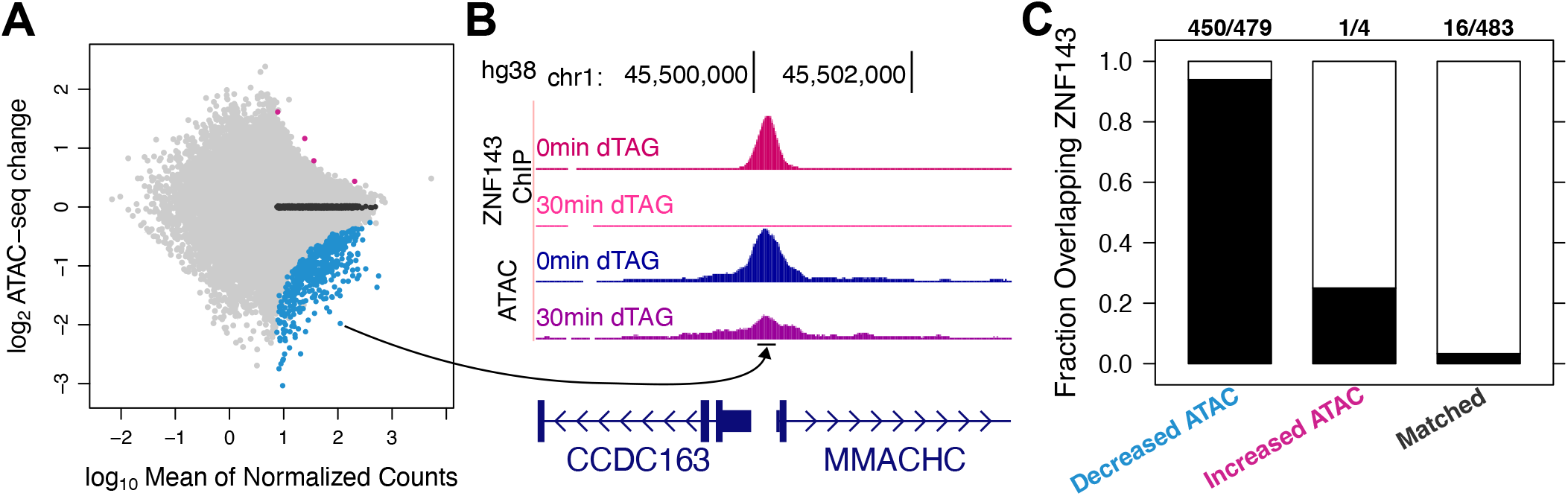
ZNF143 degradation leads to a decrease in chromatin accessibility. A) Only four ATAC-seq peaks increase chromatin accessibility after 30 minutes of ZNF143 depletion and 479 peaks decrease accessibility. B) An example ATAC-seq peak that decreases accessibility overlaps a ZNF143 binding site. C) Ninety-four percent of ATAC peaks that decrease accessibility overlap ZNF143 peaks, indicating that ZNF143 is predominantly involved in maintaining open chromatin.

### ZNF143 is an activator that predominantly regulates transcription initiation

We performed nascent RNA profiling (PRO-seq) after 30 minutes of ZNF143 depletion to identify direct ZNF143 target genes (Kwak et al. 2013). Hundreds of genes increase (*up*) and decrease (*down*) transcription after 30 minutes of ZNF143 depletion (Figure 3A). Although the expression of primary response genes goes in both directions, this does not necessarily mean that ZNF143 functions as an activator and a repressor. The transcription factors that we know most about are rapidly inducible, such as heat shock factor and nuclear hormone receptors such as the estrogen, androgen, and glucocorticoid receptors. Although genes are activated and repressed in the minutes following transcription factor induction, only the activated genes are closer to the respective binding sites of the factor (Duarte et al. 2016; Dutta et al. 2023; Guertin and Lis 2010; Guertin et al. 2014; Hasterok et al. 2023; Reddy et al. 2009). The repressed genes are no closer to the factor binding sites than unchanged control genes, so only the activated genes are considered direct *cis* targets of these transcription factors. We performed these same gene class/peak proximity analyses for the ZNF143 regulated genes and unchanged genes that are matched for expression levels (Figure 3B). Nearly all the *down* genes have their transcription start site within 500 bases of a ZNF143 binding site (Figure 3B) and the vast majority of ZNF143 binding sites are in the upstream promoter region (Figure 3C). Unlike the heat shock and hormone response systems, the opposing *up* gene class is enriched for proximal ZNF143 binding (Figure 3B), although ZNF143 binding is not limited to the promoter region (Figure 3C). These results reveal that the predominant function of ZNF143 is to activate transcription proximally and the remainder of this section will focus on determining the molecular mechanism of ZNF143-mediated activation.

**Fig. 3.**
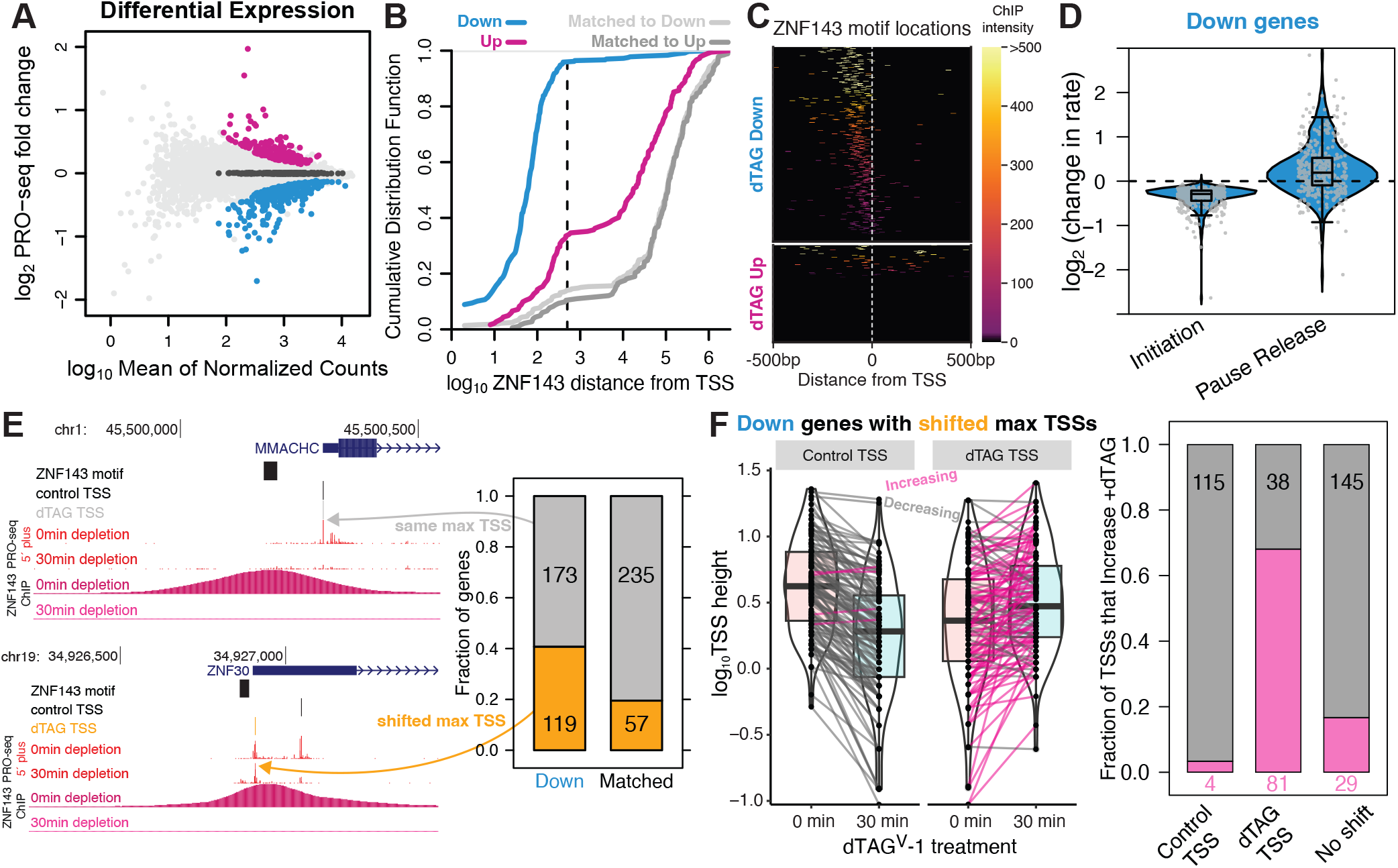
ZNF143 binds in promoters to directly regulate transcription initiation. A) Nascent transcriptomics identifies 182 genes that increase expression (*up*) and 365 genes that decrease expression (*down*) after 30 minutes of ZNF143 degradation. Genes that are matched for expression level and unchanged are in dark grey. B) Ninety-six percent of *down* genes are within 500 bases of a ZNF143 binding site. The *up* genes are also significantly closer to ZNF143 binding sites compared to genes that are matched for expression level. C) *Down* genes have ZNF143 binding in the promoter; *up* genes have no clear pattern of ZNF143 distribution and 66% have no local ZNF143 binding. D) Compartment modeling indicates that ZNF143 regulates initiation and not pause release at direct *down* target genes. Each point is a *down* gene and the y-axis values are the changes in rates that most likely explain the data. E) The transcription start sites of direct *down* target genes tend to change upon ZNF143 degradation. F) We assigned predominant transcription start sites pre/post ZNF143 degradation. The decrease in TSS usage at the control prominent TSS is accompanied by an increase in TSS usage of the prominent dTAG-treated TSS.

The transcription cycle includes many steps including chromatin decondensation, RNA polymerase recruitment/initiation, promoter-proximal pause termination or release, and elongation (Fuda et al. 2009; Wagner et al. 2023). We quantified the change in PRO-seq density within the 5′ pause region and gene bodies of *down* genes and incorporated these values into a compartment model (Figure S2) to determine whether ZNF143 predominately regulates initiation or pause release. Our modeling analysis indicates that a decrease in initiation rate at every *down* gene with ZNF143 binding within the 500 base promoter best explains the redistribution of RNA polymerase in the pause and gene body regions upon ZNF143 depletion (Figure 3D). In contrast, a comparable fraction of gene’s pause release rates change in both directions (Figure 3D). Compartment modeling of changes in PRO-seq signal cannot distinguish between decrease initiation and increasing premature pause termination because these are directly opposing rate constants (Figure S2).

To determine whether ZNF143 regulates initiation/recruitment versus premature pause termination we sought to determine whether ZNF143 affects transcription start site (TSS) usage. Our rationale is that if ZNF143 regulates initiation, then the predominant TSS would change upon ZNF143 depletion. Previous work showed that rapid depletion of the initiation factor TATA-binding protein (TBP) causes changes in TSS usage (Santana et al. 2022). The 5′ end of PRO-seq reads accumulate at a gene’s TSS and we inferred the most prominent TSS before and after 30 minutes of ZNF143 degradation from 5′ read pileups. We focused this analysis exclusively on *down* genes with ZNF143 binding within 500 bases of the TSS in the promoter region. *Down* genes with promoter bound ZNF143 shift their most prominent TSS at twice the rate compared to a control set of genes matched for TSS signal intensity (Figure 3E). For example, after ZNF143 degradation *ZNF30*’s TSS shifts from 159 to 18 bases away from ZNF143’s binding site (Figure 3E). This suggests that ZNF143’s close proximity inhibits initiation at the alternative dTAG TSS. Transcription start sites within the human genome are not focused at a single position, but occur in windows and genes can contain multiple TSS windows (Luse et al. 2020). The observed changes in the most prominent TSS could occur because signal decreases at the control TSS and an alternative less prominent TSS does not change intensity. However, we observe a coupled decrease in signal at the control TSS and an increase in signal at the ZNF143-depleted TSS (Figure 3F). This pattern suggests that not only does ZNF143 regulate transcription initiation as opposed to premature termination, but the redistribution of TSS signal suggests that ZNF143 competes with other transcription factors for RNA polymerase and/or initiation machinery. The role of ZNF143 stimulating initiation is consistent with the chromatin accessibility data from Figure 2. Although chromatin structure is influenced directly by the recruitment of chromatin remodelers and histone-modifying enzymes, the recruitment of RNA Polymerase or initiation factors can also deplete nucleosomes, thereby altering the chromatin landscape.

### ZNF143 directly represses genes by binding directly over transcription start sites

The canonical molecular functions of ZNF143 are succinctly described as DNA binding and stimulation of transcription initiation. However, it is not immediately clear how these functions lead to the direct *cis* repression of ZNF143 target genes observed in Figure 3B. ZNF143 binds both upstream and downstream of *up* genes at comparable frequencies (Figure 3C), but it directly binds over the TSS of five genes (Figure 4A&B). ZNF143 exhibits stronger binding over the TSS of four out of these five genes compared to matched and *down* genes with TSS-bound ZNF143 (Figure 4B). These observations support a model in which ZNF143 binding directly over the TSS competes with RNA Polymerase for access to the initiator sequence. The distribution of 5′ PRO-seq reads remains consistent with or without ZNF143 depletion at the one *down* gene, *FIS1*, that exhibits weak ZNF143 binding over the TSS (Figure 4C). However, a closer inspection of this gene reveals that the most prominent TSS changes upon ZNF143 depletion and 5′ PRO-seq signal substantially increases at this new TSS (Figure 4D). A much stronger ZNF143 binding site is located directly downstream of this *FIS1* TSS-isoform (Figure 4D). This observation suggests an independent repressive mechanism, where ZNF143 binding directly downstream of a TSS acts as a molecular roadblock, which will inhibit RNA Polymerase from progressing into the gene body and effectively inhibit initiation of new RNA Polymerases.

**Fig. 4.**
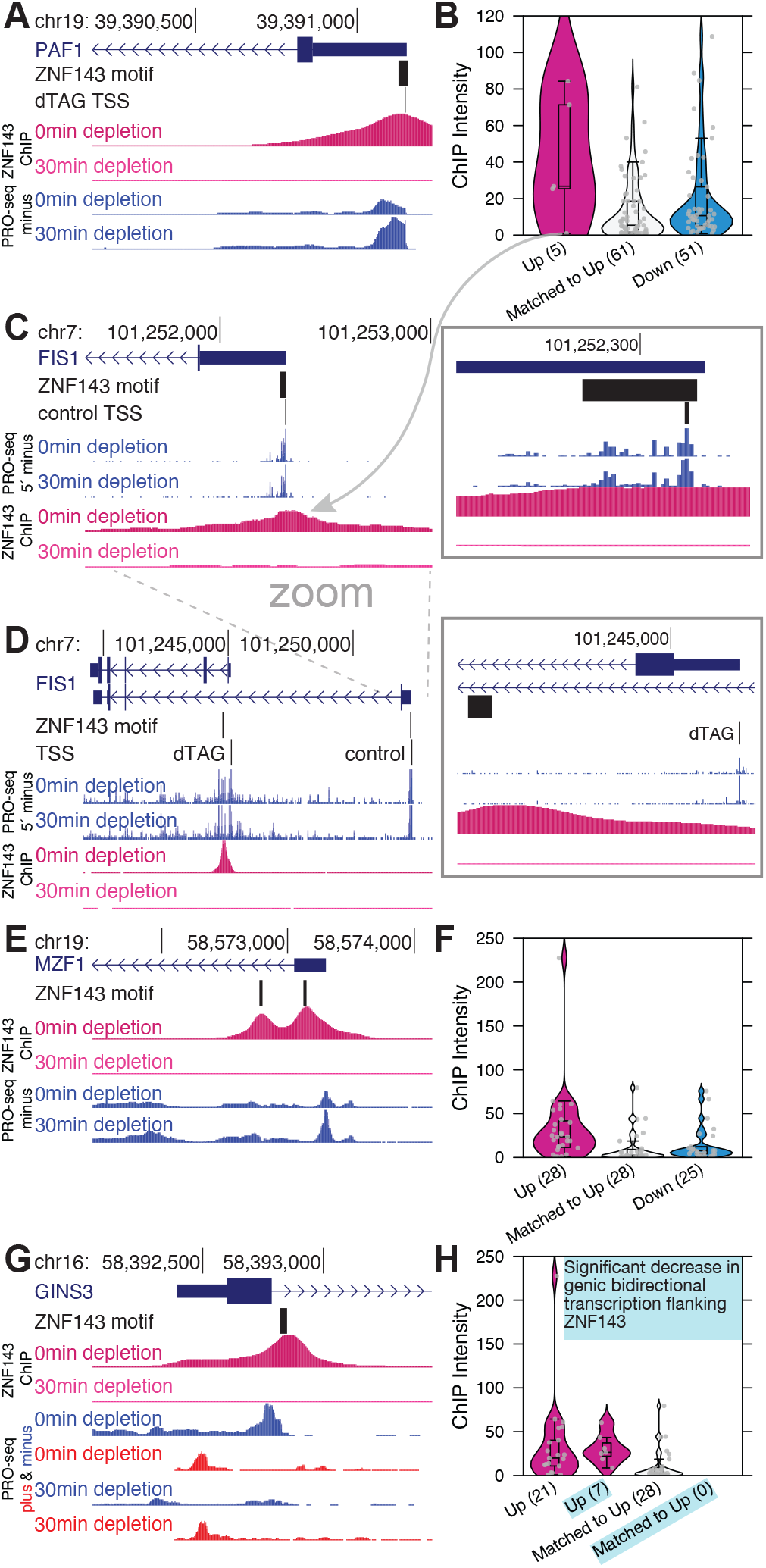
ZNF143 occludes sites of initiation and acts as a molecular roadblock to directly repress genes in *cis*. A) ZNF143 binds a motif that spans *PAF1*’s transcription start site. *PAF1* transcription increases upon ZNF143 depletion. B) ZNF143 binds strongly over the TSS of 4/5 genes that increase transcription upon ZNF143 depletion. C) *FIS1* is the *up* gene from panel B with weak TSS binding of ZNF143 and no discernible changes in TSS usage occur upon depletion, as measured by PRO-seq 5′ end pile ups. D) ZNF143 binds directly downstream from the TSS of a different isoform of *FIS1*. This *FIS1* isoform becomes the most prominent *FIS1* TSS upon ZNF143 degradation. E&F) ZNF143 degradation from strong binding sites directly downstream of transcription start sites is associated with *up* genes. G&H) Genic bidirectional transcription at ZNF143 binding sites downstream of *up* genes significantly decreases at 7/28 *up* genes.

### ZNF143 directly represses genes by acting as a molecular roadblock immediately downstream of transcription start sites

*FIS1* was not initially classified as a ZNF143-downstream gene because we inferred a single isoform of each primary transcript using the control data (Anderson et al. 2020). We then extended our analysis to the twenty-eight *up* genes with ZNF143 binding within 500 bases downstream of their control TSSs (Figure 4E). These 28 genes have strong ZNF143 binding downstream of their start sites compared to matched and *down* genes with downstream ZNF143 peaks (Figure 4F). These results are consistent with a molecular roadblock model, whereby strong transcription factor binding can inhibit RNA polymerase during early elongation when it is accelerating and more vulnerable to pausing, backtracking, and disassociation. Beyond ZNF143’s role in DNA binding, we have established that it also stimulates initiation. Importantly, RNA polymerase initiates transcription at bidirectional regions across the genome, not solely at genic TSSs (Core et al. 2014, 2008). Bidirectional transcription (Azofeifa and Dowell 2017; Wang et al. 2019), especially an RNA polymerase colliding head on with a genic preinitiation complex or paused RNA polymerase, could also act to repress transcription. We find that 7 out of these 28 downstream ZNF143-binding genes have significantly reduced bidirectional transcription after ZNF143 depletion (Figure 4G&H). This evidence supports the hypothesis that ZNF143 not only activates but may also repress transcription by promoting RNA polymerase initiation, highlighting the importance of proposing specific mechanisms when claiming transcription factors have dual activation/repression functions.

### ZNF143 occludes downstream sequence elements to repress transcription

ZNF143 binds to the promoters of three *up* genes that have an upstream shift in their TSS after ZNF143 depletion (Figures 5B&S4). We hypothesized that ZNF143 binding occludes a downstream sequence motif that contributes to initiation. The downstream promoter element (DPE) located +28 to +32 downstream of TSSs was the first downstream element shown to contribute to initiation by interacting with TFIID (Burke and Kadonaga 1997). ZNF143 blocks a DPE that is located at positions +27 to +31 within the *LYSMD1* gene. Additionally, YY1, an initiation factor (Athanikar et al. 2004; Seto et al. 1991), has a binding motif that is commonly found directly downstream of TSSs and between +100 to +300 positions downstream (Benner et al. 2013; Dudnyk et al. 2024) (Figure 5A). ZNF143 binds upstream of *LYSMD1, GMPR2*, and *ZNF583*, and upon ZNF143 depletion, a YY1 motif becomes accessible at each of these genes (Figures 5B&S4). YY1 binding at the exposed site may facilitate efficient initiation at the upstream TSS. This YY1 occlusion mechanism may be acting simultaneously as a roadblock and/or directing genic bidirectional transcription for the upstream dTAG TSS (Figure S4). These three *up* genes are unique because ZNF143 is promoter-bound and ZNF143 depletion causes an upstream TSS shift. This mechanism cannot explain repression at *up* genes with promoter-bound ZNF143 and no upstream TSS shift upon depletion.

**Fig. 5.**
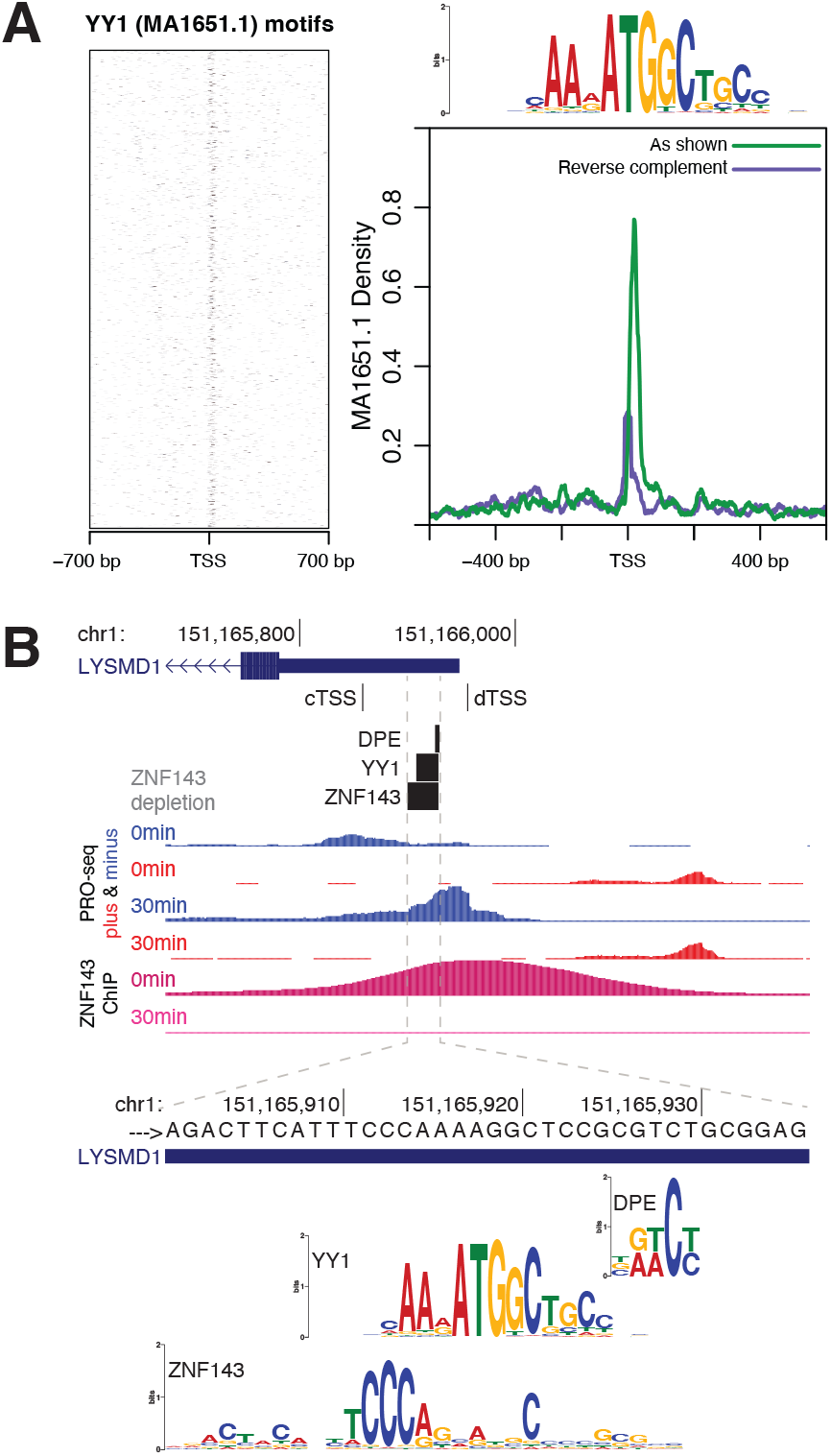
ZNF143 occludes downstream sequence elements to repress transcription. A) Transcription factor YY1’s motif (MA1651.1) is enriched directly downstream of TSSs. B) ZNF143 depletion reveals an accessible YY1 and DPE that allows usage of a TSS that is 28 bases upstream of the ZNF143 binding site in the *LYSMD1* promoter (cTSS: control max TSS position. dTSS: dTAG max TSS position).

### SP1 redistributes to ZNF143 binding sites to stimulate initiation

ZNF143 bound the promoters of 25 *up* genes with distances and intensities comparable to those of *down* genes and no TSS shift upstream of ZNF143’s binding site (Figure 3C). Given the limited number of *up* genes, we catalogued all bidirectionally transcribed putative regulatory elements across the genome and analyzed transcriptional changes in these regions after ZNF143 depletion (Love et al. 2014; Wang et al. 2019) (Figure 6A&B). We then carried out *de novo* motif discovery within the bidirectionally transcribed regions of both *down* and *up* classes. As anticipated, the ZNF143 motif was prevalent in the *down* class; however, its presence in the *up* class was unexpected (Figure 6B). Additionally, the SP transcription factor family DNA motif was identified *de novo* in the *up* class (Figure 6B). Ninety-nine bidirectional *up* regions have both ZNF143 ChIP peaks and ZNF143 motifs, with nineteen of the ZNF143 motifs overlapping SP motifs (Figure 6C&D). We performed SP1 ChIP-seq before and after 30 minutes of ZNF143 depletion to address the possibility that SP1 is redistributing to these sites when ZNF143 vacates. Each of the these 19 regions overlapped an SP1 ChIP-seq peak and 13 of the peaks increased SP1 intensity upon ZNF143 depletion (Figure 6E). SP1 also regulates initiation (Dutta et al. 2023; Gill et al. 1994), so these results are consistent with a model whereby an canonical activator can repress a gene if it competes with a more potent activator for the same stretch of DNA. SP1 is clearly not replacing ZNF143 at all nineteen sites, but recall that ZNF143 binds a 29 base region and the ZNF143 motif may be overlapping the motifs for other initiation factors.

**Fig. 6.**
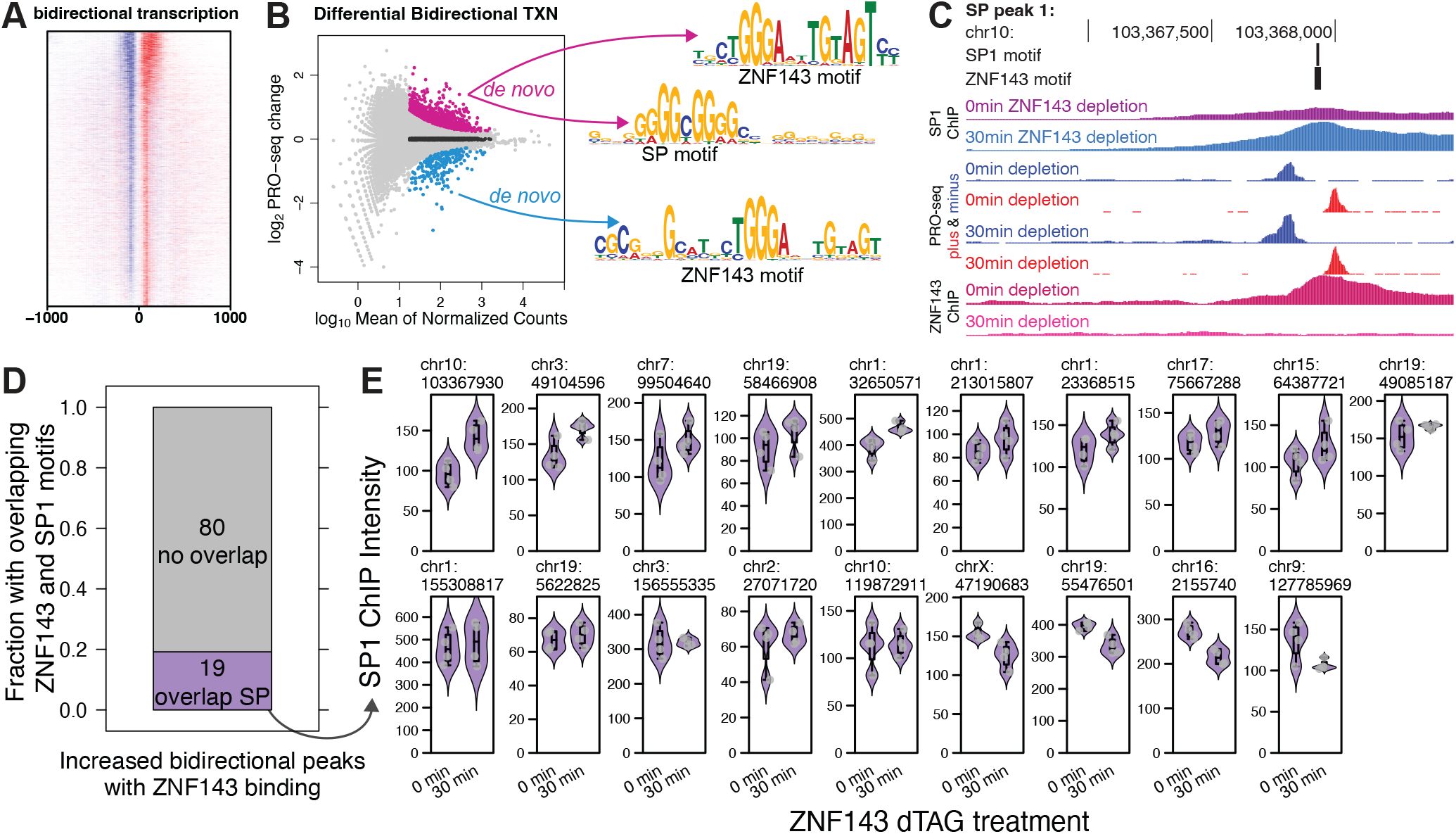
ZNF143 may compete with SP1 at overlapping binding sites to facilitate transcription initiation. A) We called sites of bidirectional transcription genome-wide and aligned on their center position (Wang et al. 2019). B) We identified the SP factor family of motifs *de novo* exclusively in bidirectionally transcribed that increase transcription. The ZNF143 motif was identified *de novo* in both *up* and *down* birdirectionally transcribed regions. C) SP1 binding intensity, as measured by ChIP-seq, increases at a bidirectionally transcribed region that has overlapping SP1 and ZNF143 motifs. D) Nineteen of the bidirectional transcription regions that increase and have ZNF143 binding have SP1 binding and the SP1 sequence motifs that overlap the ZNF143 sequence motifs. E) Thirteen of the nineteen regions from panel (D) increase SP1 binding upon ZNF143 degradation.

SP1 motifs were among the first promoter sequence elements to be identified as regulatory (McKnight and Kingsbury 1982) and SP1 has had a defined role in initiation for thirty years (Gill et al. 1994). Recent studies have further refined SP motifs as defining elements of promoter architecture in the genome (Benner et al. 2013; Dudnyk et al. 2024; Jones et al. 2024). Motifs recognized by paralogs of ZNF143, along with NRF1, ETS, NFY, CREB/ATF (bZIP), and SP/KLF factors, are considered critical in dictating promoter structure and initiating transcription (Benner et al. 2013; Dudnyk et al. 2024; Jones et al. 2024). The sequence-specific transcription factors that bind these motifs may be a class of sequence-specific transcription factors that recruit the initiation machinery and RNA polymerase. Our findings confirm the enrichment of these motifs within bidirectional promoters across the human genome (Figures 7A&S3). Sixteen *up* genes with ZNF143 in the promoter have an SP motif within an SP1 ChIP-seq peak that overlaps the ZNF143 element and 10 of the 16 have an increase in SP1 ChIP signal after ZNF143 depletion (Figure 7B). The promoters of *ZNF688, UTP3*, and *PYCR2* exhibit a statistically significantly (FDR < 0.01) increase in SP1 signal upon ZNF143 depletion; however, the redistribution of SP1 at other genes is either more subtle or non-existent (Figure 7B). We searched for the motifs of these sequence-specific initiation transcription factors within the 29mer ZNF143 motif of the 28 genes with promoter-bound ZNF143. At least one of the bZIP, SP/KLF, ETS, and NRF1 motifs overlapped the ZNF143 motif (Figure 7C&D) in all genes except *TSNAXIP1* (Figure 7D). We propose that other activating factors replace ZNF143 at these promoters when ZNF143 is depleted. For example, ZNF143 is ablated and SP1 intensity reduced two-fold at their overlapping binding site in the promoter of *GSK3A*, but we hypothesize that this reduction of binding enables an ETS factor to bind strongly and stimulate transcription (Figure 7D). We also observe the same overlap of ZNF143 and NRF1/ETS/SP motifs at bidirectionally transcribed regions from Figure 6D (Figure 7E). These results suggest that a canonical activator may repress genes by competing for DNA binding with a more potent activator. However, we appreciate that the molecular calculus for such a mechanism is complex. For instance, a very strong activator might have a brief residency time on DNA, making its overall contribution minimal, while a modest activator could have more stable DNA binding and effectively stimulate transcription. Given that each promoter has a unique sequence context, predicting transcriptional outcomes solely based on sequence is challenging. To fully understand the competitive interactions between two transcription factors targeting the same DNA region, one must consider: 1) their relative binding efficiencies and residency times within specific chromatin contexts; 2) the transcription cycle stages each factor influences; 3) the relative activation potency of each factor; and 4) which steps in the transcription cycle limit the output of target genes.

**Fig. 7.**
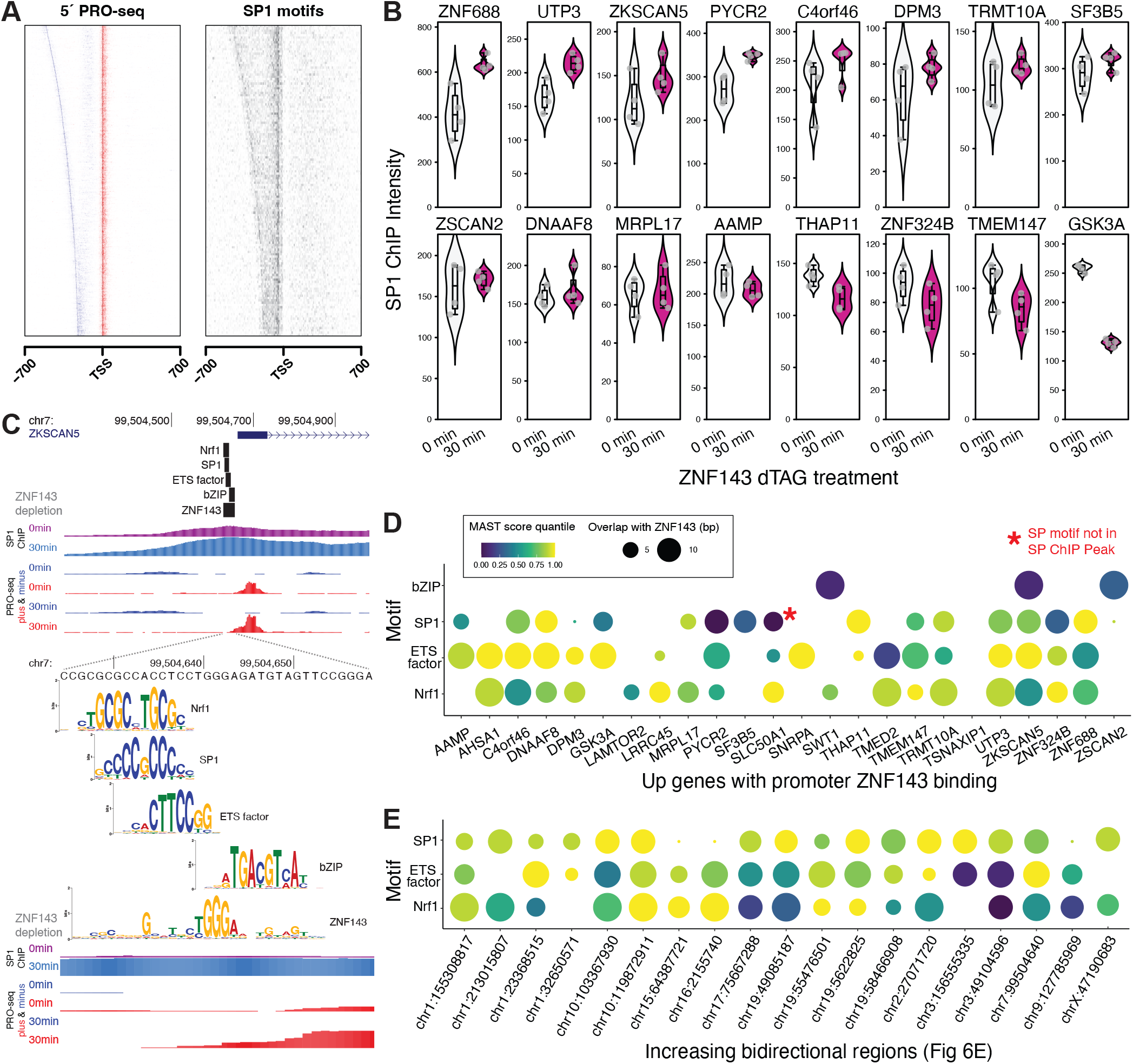
ZNF143 may compete with other promoter-bound sequence-specific transcription factors to regulate transcription. A) We identified promoter regions within sense and divergent TSSs called from pileups of the 5’ ends of PRO-seq reads. SP motifs identified by FIMO (Grant et al. 2011) are enriched in regions between bidirectional start sites. B) After ZNF143 degradation, sixteen genes that increased transcription also had an SP1 ChIP-seq peak and motif overlapping the ZNF143 motif. The corresponding SP1 binding intensity (ChIP) increased for 10 of these genes. C) Other promoter-enriched motifs (Nrf1, bZIP, and ETS) also overlapped ZNF143 motifs in the promoter of *ZSCAN5*. D) Other motifs that overlap ZNF143 motifs in promoters of *up* genes have varying levels of motif conformity and overlap. The red asterisk marks an SP motif that overlaps ZNF143 but is outside of an SP1 ChIP peak. Circle size represents amount of overlap with the 29 base ZNF143 motif and circle color represents motif conformity as measured by MAST score quantiles that were calculated by querying the weight matrices against 1 million random sequences. E) The bidirectional regions identified in Figure 6E also have other promoter-enriched motifs that overlap ZNF143.

## Discussion

When discussing the context-specific dual roles of transcription factors as both activators and repressors, it is crucial to propose an underlying mechanism (Figure 8). The most well characterized example of dual activator/repressor function in gene regulation is that of the *λ* repressor. The *λ* repressor is an activator that interacts with the bacterial RNA polymerase to stimulate initiation, but it was named *repressor* because it represses transcription of the lytic genes, which account for nearly all the phage genes. The *λ* repressor binds to a precise position in its own promoter to stimulate RNA polymerase initiation and activate the gene encoding itself (Ptashne 2004). This same binding event blocks RNA polymerase from accessing and initiating transcription of lytic genes (Johnson et al. 1979; Meyer et al. 1980; Ptashne 2004). The molecular functions of both *λ* repressor and ZNF143 are to bind DNA and stimulate initiation. Binding of ZNF143 to DNA is necessary to stimulate initiation; like *λ* repressor, ZNF143 typically binds upstream of a initiation site to activate transcription. Binding in a promoter and stimulating initiation may result in repression if ZNF143 displaces a more potent stimulator of initiation. Binding over an initiator sequence or within a gene body can cause repression via multiple described mechanisms. While ZNF143 can both activate and repress genes, its molecular functions of DNA binding and promoting initiation remain constant.

**Fig. 8.**
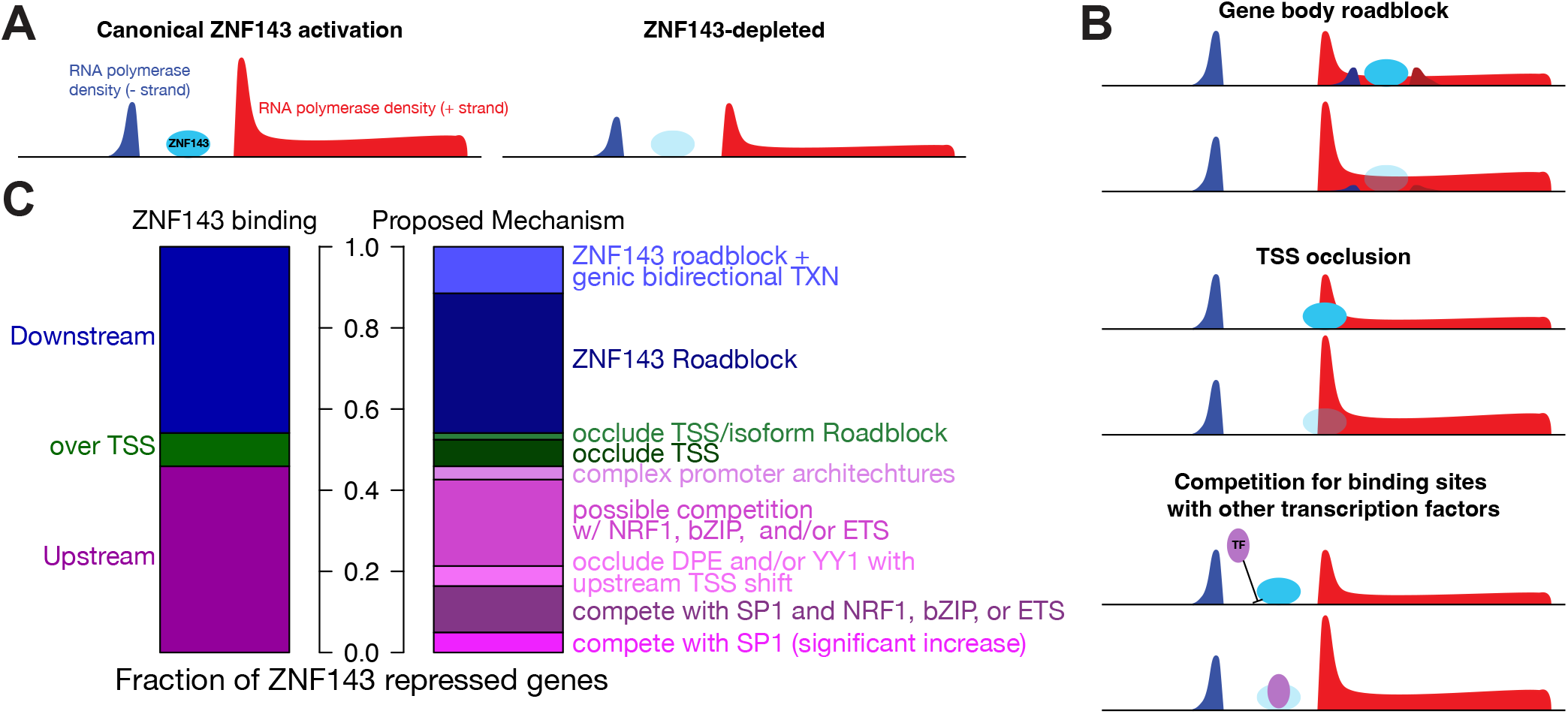
The molecular function of ZNF143 is to bind DNA and facilitate initiation. A) ZNF143 binds to the promoter and stimulates transcription at the majority of its target genes. B) ZNF143 binding can cause repression if binding occurs over a TSS, immediately downstream of a TSS, or in competition with other activators. C) ZNF143’s repression mechanisms can be further specified depending on the local environment of the ZNF143 binding site. The number of genes in the left barchart from top to bottom is 28, 5, and 28. The right bar chart top to bottom has the following number of genes in each category: 7 (roadblock/bidirectional), 21 (roadblock), 1 (roadblock and TSS occlusion), 4 (TSS occlusion), 2 (complex promoters with no proposed mechanism), 13 (possible competition with sequence-specific promoter factors), 3 (occlude DPE and/or YY1), 7 (competition with SP1 and another factor), and 3 (competition with SP1).

Many transcription factors are described as having both activator and repressor activities and *context specificity* is often vaguely invoked as the explanation. There are conflicting reports on whether MYC acts as an activator at specific genes, an amplifier of all genes, or a direct repressor at some genes (Lin et al. 2012; Lorenzin et al. 2016; Nie et al. 2012; Sabo et al. 2014; Walz et al. 2014). The acute depletion of an auxin inducible degron tagged MYC within 30 minutes, coupled with nascent RNA sequencing, allowed for the identification of primary MYC-responsive genes (Muhar et al. 2018). 98% of these genes were repressed, indicating that the direct effect of MYC regulation is transcriptional activation of only a fraction of the expressed genes in a cell (Muhar et al. 2018). OCT4 is one of the four pluripotency factors, the expression of which is sufficient to reprogram differentiated fibroblasts into induced pluripotent stem cells (Takahashi and Yamanaka 2006). However, the genes OCT4 regulates in pluripotent stem cells are difficult to identify, as the half-life of its protein and mRNA are much too long for traditional knockdown methods to isolate the primary effects of depletion (Bates et al. 2021). A recent study compared extended knockdown to rapid depletion with targeted protein degradation and found that only the latter was able to identify that the primary effect of OCT4 on transcription is the activation of pluripotency factors and that the delayed activation of trophoblast-associated genes is a secondary effect of OCT4 depletion (Bates et al. 2021). A key takeaway from these studies is that these factors directly activate transcription of their target genes. The growing list of transcription factors that can be acutely perturbed provides evidence that most factors do not activate some direct targets and repress others. Another theme is that, in contrast to extended knockdown, acute depletion of most sequencespecific factors affects transcription of a limited number of primary response genes (Sheppard et al. 2021; Stengel et al. 2021).

There is substantial data suggesting that transcription factors specialize and recruit either corepressors or coactivators. We can conceive of exceptions to this rule, but we would argue that a factor can become a fundamentally different protein with distinct functions based on ligand binding status or post-translational modifications. The most well-characterized example of this is the thyroid hormone receptor (THR). THR binds to DNA and recruits corepressors in the absence of thyroid hormone (Hörlein et al. 1995); THR recruits coactivators when bound by thyroid hormone (Chen et al. 1997; Fondell et al. 1996; Grøntved et al. 2015; Lin et al. 1997). Post-translational modifications, such as phosphorylation or acetylation, fundamentally change the identity of a protein and this may result in the ability to differentially interact with cofactors. Although we were unable to identify well-characterized examples of posttranslational modifications that switch interaction partners from coactivators to corepressors or vice versa, this mechanism has been suggested for STAT3, which recruits corepressors exclusively when acetylated (Gambi et al. 2019). Extending this general mechanism, a transcription factor may be bound to DNA, but only interact with coactivators when posttranslationally modified. In the absence of the modification, the factor would be a functional repressor by occluding the binding sites of other activating factors and failing to recruit coactivators.

Another key part of context for defining the role of a TF is promoter strength. Transcription factor activity modeled in *E. coli* demonstrates that a TF can appear to change from an activator and repressor or vice versa depending on promoter strength and which step of the transcription cycle is regulated by the TF (Ali et al. 2023). In their model, transcription factors were characterized by their ability to either regulate the stability of RNA polymerase or promote transcription initiation. They found that TFs with strong stabilizing functions but weaker initiating abilities moved from activation to repression as promoter strength increased. We can envision how the same RNA polymerase stabilization mechanism might appear to repress transcription in the presence of a strong promoter if the stabilization interferes with the ability of RNA polymerase to begin transcription. On the other end of the spectrum, TFs that were strong initiators but weaker stabilizers moved from repression to activation with increasing promoter strength. Once again, we see that a TF appears to change activity when in reality the strong promoter may make it easier for transcription to initiate when perhaps other factors are able to improve RNA polymerase stability. The importance of promoter strength in eukaryotes is highlighted by an effector domain mutagenesis and mapping study that identified effector domains with the ability to seemingly both activate and repress transcription (DelRosso et al. 2023). Here the definition of “activator” and “repressor” becomes important, as this study defined activators as effector domains able to increase expression as compared to basal levels from a minimally active promoter and repressors as those able to decrease expression as compared to basal levels from a constitutively active promoter. According to these definitions, it is possible that a “bifunctional” effector domain is actually a weak activator if it promotes transcription, but at a lower level than the constitutively active promoter. Although the details of transcription regulation in prokaryotes differs from eukaryotes, the overall principle appears to be similar in that promoter strength, combined with the particular role of a TF, both contribute to whether a particular TF appears to be activating or repressing transcription. Understanding the relationship between promoter strength and the specific functions of TFs can explain their seemingly dual role in activating and repressing gene expression.

Although this study convincingly demonstrates that ZNF143 depletion can lead to the repression of local genes via our proposed plausible mechanisms (Figure 8), we have yet to distinguish between incidental regulation and conserved evolutionary mechanisms that specifically govern these genes. The identity of the ZNF143-repressed genes suggest some coordinated regulatory control. *FIS1* is a mitochondrial fission gene and ZNF143 positively regulates nuclearly encoded mitochondrial genes (Magnitov et al. 2024). We speculate that activating nuclearly encoded mitochondrial genes that are involved in mitochondrial biogenesis and function while simultaneously repressing fission genes can be a strategic response to enhance mitochondrial function and biogenesis. This coordinated response may build a robust mitochondrial network to meet increased metabolic demands or recover from damage more effectively. Another gene that is repressed in *cis* by ZNF143 is *THAP11. THAP11* binds to a nearly identical sequence motif as ZNF143 (Ngondo-Mbongo et al. 2013; Vinckevicius et al. 2015), although there is no clear evolutionary conservation of their respective DNA binding domains. The regulation of *THAP11* by ZNF143 is not clear from our data, although we propose that ZNF143 displaces an ETS factor (Figure 7). Regulation of *THAP11* is likely more complicated, as close inspection of the locus reveals an unannotated TSS that is flanked by two ZNF143 ChIP peaks, which are also likely THAP11 binding sites (Figure S5). Moreover, usage of this unannotated TSS increases substantially upon ZNF143 degradation. Given the interplay between ZNF143 and the regulation of genes like *FIS1* and *THAP11*, it is likely that at least some aspects of regulatory repression are crucial for maintaining homeostasis and orchestrating cellular responses to metabolic changes.

This work underscores the significance of using rapidly inducible systems to study molecular mechanisms. Additionally, we highlight that while genomic experiments and analyses indicated ZNF143 was repressing a subset of genes in *cis*, a more detailed examination of these repressed genes was essential to suggest mechanisms of repression. We recognize that a thorough understanding of biological systems necessitates an in-depth grasp of the individual components and mechanisms. We therefore anticipate that the pendulum of scientific inquiry will swing back from broad molecular genomics studies, high throughput reporter assays, and large scale screens towards more focused, mechanistic investigations.

## Methods

### HEK-293T culture and ZNF143-dTAG clone generation

HEK293T cells were cultured at 37°C with 5% carbon dioxide in DMEM media supplemented with 10% fetal bovine serum (FBS), 100 U/mL penicillin-streptomycin, 2.2mM L-glutamine, and 1mM sodium pyruvate. We endogenously tagged ZNF143 at the C-terminus in HEK-293T cells as previously described (Sathyan et al. 2020; Scott et al. 2024). We targeted ZNF143 endogenously using CRISPR loaded with the sgRNA (GAGGATTAATCATC-CAACCCTGG). We cleaved the hSpCas9 plasmid PX458 (Addgene #48138) with the enzyme BbsI, then annealed oligonucleotides 5’-CACCGAGGATTAATCATCCAACCC-3’ and 5’-AAACGGGTTGGATGATTAATCCTC-3’, and inserted the annealed product into the plasmid. We generated a linear homology-directed repair donor by amplifying the pCRIS-PITChv2-dTAG-Puro plasmid (Addgene #91796) with the primers in Table 1 (Nabet et al. 2018). Following transfection of the donor DNA and Cas9/sgRNA plasmid into HEK-293T cells, cells were selected and cloned as described in (Sathyan et al. 2020). We selected cells in media with 1 µg/ml puromycin and confirmed successful integration by Western blot as previously described (Scott et al. 2024). After obtaining a correctly-tagged clone (ZD29) we passaged cells thawed from frozen aliquots until the desired number of cells for each experiment was reached, and then treated with dTAG^V^-1 and collected cells for ATAC-seq, PRO-seq, and ChIP-seq.

**Table 1.**
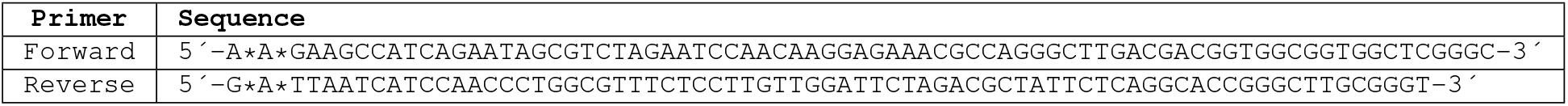
Homology directed repair primers. These primers amplify FKBP^F36V^-2xHA-P2A-Puro with 50 bases of ZNF143 homology. The asterisks mark phosphorothioate bond modifications to minimize degradation.

### Genome browser visualization

Genome browser (Kent et al. 2002) images were taken from the following track hub: http://guertinlab.cam.uchc.edu/znf143_hub/hub.txt.

### ATAC-seq library preparation

We prepared ATAC-seq libraries as previously described (Grandi et al. 2022; Scott et al. 2024). We treated cells with either DMSO or dTAG^V^-1 in DMSO for thirty minutes. After treatment, we aspirated the media and scraped the cells on ice and centrifuged them at 500g for 5 min at 4°C. We resuspended cell pellets in lysis buffer (0.1% NP40, 0.1% Tween-20, 0.1% digitonin) made in cold resuspension buffer (10mM Tris-HCl ph 7.5, 10mM NaCl, 3mM MgCl_2_ in water) and then incubated on ice for 3 minutes. We mixed lysates with wash buffer (0.1% Tween-20 in resuspension buffer) and centrifuged cells at 500g for 10 min at 4°C. We resuspended the pellets in a transposition mixture (1x TD buffer, 0.1% ultrapure distilled water, 0.01% digitonin, 0.1% Tween-20 in PBS) and added 2.5µl of TDE1 Tn5 transposase (Illumina Tagment DNA TDE1 Enzyme and Buffer Kit). We incubated the transposition reaction at 37°C for 30 min. We isolated DNA from the reaction with the DNA Clean and Concentrator-5 kit. We added adapters and amplified DNA over 8 cycles of PCR using the NEBNext Ultra II kit. We purified and size selected the library by incubating samples with AMPure XP beads (1.8x buffer to sample ratio) and eluting with nuclease-free water.

### ATAC-seq analyses

We aligned raw sequence data to the *hg38* genome assembly with `bowtie2`, converted to *BAM* format with `samtools`, and then to *bigWig* format with `seqOutBias` (Langmead and Salzberg 2012; Li et al. 2009; Martins et al. 2018). Peak calling with MACS3 (Zhang et al. 2008) employed the following arguments: `-q 0.01 -keep-dup all -nomodel -shift -100 -extsize 200`. We counted reads in peaks with the `bigWig` R package and called differentially accessible regions with `DESeq2` (Love et al. 2014; Martins 2014).

### PRO-seq library preparation

We prepared PRO-seq libraries as previously described (Mahat et al. 2016; Sathyan et al. 2019). After thirty minutes of treatment with either 500nM dTAG^V^-1 or DMSO, we washed cells ice cold PBS. We collected cells by adding buffer W (10mM Tris-HCl pH 7.5, 10mM KCl, 250mM sucrose, 5mM MgCl_2_, 1mM EGTA, 10% glycerol, 0.5mM DTT, 0.004U/mL SUPERaseIN RNase inhibitor, and fresh protease inhibitors) and scraping the cells, followed by centrifugation at 500g for 5 minutes and resuspension in buffer W. We added buffer P (10mM Tris-HCl pH 7.5, 10mM KCl, 250mM sucrose, 5mM MgCl_2_, 1mM EGTA, 0.05% Tween-20, 0.1% NP40, 10% glycerol, 0.5mM DTT, 0.004U/mL SUPERaseIN RNase inhibitor, fresh protease inhibitors) for 5 minutes to permeabilize the cells. We centrifuged and resuspended the cells in buffer W twice before pelleting the cells again and resuspending in buffer F (50mM Tris-HCl pH 8, 5mM MgCl_2_, 1.1mM EDTA, 40% Glycerol and 0.5mM DTT). We snap froze aliquots in liquid nitrogen and kept them stored at -80°C. PRO-seq library prep was done in a method based on previously described protocols (Judd et al. 2020; Mahat et al. 2016). After the run-on reaction, we added adapters that included a random eight base unique molecular identifier (UMI) on the 5′ end of adapter that is ligated to the 3′ end of the nascent RNA. We eluted and reverse transcribed the RNA and performed 10 cycles of PCR. We purified the PCR reactions with a MinElute PCR purification kit and did not perform size selection in an effort to preserve short nascent RNAs in our libraries.

### PRO-seq analyses

Quality control and read alignment were performed as described previously (Scott et al. 2022). We used cutadapt to remove adapters from our reads (Martin 2011), and `fqdedup` to deduplicate our libraries with the 3′ UMIs (Martins and Guertin 2018). We removed 8-mer UMIs and reverse complemented the reads with `FASTX-Toolkit` (Gordon 2010). We aligned to *hg38* with `bowtie2` (Langmead and Salzberg 2012), sorted reads with `samtools` (Li et al. 2009), and used `seqOutBias` (Martins et al. 2018) to convert reads to *bigWig* files. We used `primaryTranscriptAnnotation`and `TSSinference` to infer gene annotations from our PRO-seq data (Anderson et al. 2020; Dong and Guertin 2024). We quantified gene expression by querying the *bigWig* files within the gene annotation coordinates with the `bigWig` R package and UCSC Genome Browser Utilities (Kent et al. 2010; Martins 2014). We found differentially expressed genes with `DESeq2` (Love et al. 2014). We identified regions of bidirectional transcription with `dREG` (Wang et al. 2019). We identified overrepresented motifs *de novo* in dREG-defined regions with MEME (Bailey et al. 2015b). We modeled the rates of transcription initiation and pause release using a compartment model as previously described (Dutta et al. 2023).

### ChIP and library preparation

We fixed cells with 1% formaldehyde for 10 minutes at 37°C and quenched them with 125mM glycine for 10 minutes at 37°C. We then moved plates to ice and washed and scraped the cells into ice-cold PBS containing fresh protease inhibitors. We centrifuged cells in aliquots of 2x10^7^ cells at 1500g for 5 minutes, snap froze them in liquid nitrogen, and stored them at -80°C. After thawing the pellets, we lysed the cells in 1mL lysis buffer with protease inhibitors (0.5% SDS, 10mM EDTA, 50mM Tris-HCL pH 8.0) on ice for 10 minutes. Lysates were sonicated at 70% amplitude for 15 seconds on and 45 seconds off for 4 sets of 20-minute cycles. We moved sonicated lysates to 1.5ml tubes and clarified by centrifugation at 14,000rpm for 10 min in 4°C. We diluted 50µL of the supernatant into 760µL ChIP Dilution Buffer (0.01% SDS, 1.1% Triton X-100, 1.2mM EDTA, 167mM NaCl, 16.6mM Tris-HCl pH 8.0, fresh protease inhibitors) for a total of 1x10^6^ cells per replicate. 1ml (4x10^6^ cells) was aliquoted into each of 3 tubes with antibody (2µg anti-HA Invitrogen #26183, 8µg anti-SP1 Santa Cruz Biotechnology sc-17824 X, or mock IP) and incubated with end-over-end rotation at 4°C overnight. We washed 80µL Protein A/G Magnetic Beads (New England Biolabs) per sample with bead washing buffer (PBS with 0.1% BSA and 2mM EDTA) prior to incubating with samples for 90 minutes with rotation at 4°C. After incubation the samples were washed once each with low salt immune complex buffer (0.1% SDS, 1% Triton x-100, 2mM EDTA, 150mM NaCl, 20mM Tris HCl pH 8.0), high salt immune complex buffer (0.1% SDS, 1% Triton x-100, 2mM EDTA, 500mM NaCl, 20mM Tris Hcl pH 8.0), LiCl immune complex buffer (0.25M LiCl, 1% Igepal, 1% sodium deoxycholate, 1mM EDTA, 10mM Tris-HCl pH8.0), and 1xTE (10mM Tris-HCl, 1mM EDTA pH 8.0). We eluted the immune complexes in elution buffer (1% SDS, 0.1M sodium bicarbonate). We incubated each sample with 1µl RNase A for 10 min at 37°C. We digested the proteins by adding 5µl Proteinase K and incubating the samples in a 65°C water bath overnight. We purified DNA with a MinElute PCR purification kit, and prepared libraries with a NEBNext Ultra II Library Prep Kit.

### ChIP analysis

We removed adapters with `cutadapt` (Martin 2011) and aligned to the *hg38* genome assembly with `bowtie2` (Langmead and Salzberg 2012). We sorted aligned reads with `samtools` and used seqOutBias to generate `bigWig` files (Li et al. 2009; Martins et al. 2018). We called peaks with macs3 (Zhang et al. 2008). We quantified peak intensities by querying `bigWig` files in 400 base pair windows centered on peak summits and normalized the counts with `DESeq2` size factors (SP1 ChIP) or read depth-calculated size factors (ZNF143 ChIP) (Love et al. 2014). We found *de novo* motifs with `MEME` (Bailey et al. 2015b) and later `STREME` (Bailey 2021) through rounds of iterative motif analysis whereby we used `MAST` (Bailey and Grib-skov 1998) to identify and remove the most common motifs for SP1 or ZNF143 until we stopped finding significant *de novo* motifs.

### Data Access

All analysis details and code are available at https://github.com/guertinlab/znf143_degron. Raw sequencing files and processed *bigWig* files are available from GEO accession record GSE266491 (PRO-seq), GSE266490 (ATAC-seq), and GSE266489 (ChIP-seq).

## ACKNOWLEDGEMENTS

This work was funded by R35-GM128635 to MJG. We thank the 2023 Cold Spring Harbor Laboratory Gene Expression course participants: Rafiou Agoro, Anusha Bhatt, Nicole Brossier, Magda Bujnowska, Terri Cain, Frederic Carew, Mariano Colon-Caraballo, Nadia Côté, Elisa Gorostieta Salas, Rebecca Mello, Erik Rodriguez Teran, Sanket Shah, Jenny Shim, Tejus Sreelal, Bradley Stevens, and Dejauwne Young. The CSHL GeneX participants each generated a replicate of the ATAC-seq data under the direction of TGS and MJG. We also thank Sathyan Kizhakke Mattada for generating the degron-tagged cell line and PRO-seq libraries.

## AUTHOR CONTRIBUTIONS

JD, TGS, RM, and MJG analyzed the data. JD, TGS, and MJG performed the experiments. JD, TGS, and MJG conceptualized and developed the project. JD, TGS, and MJG wrote the manuscript.

---

**Fig. S1.**
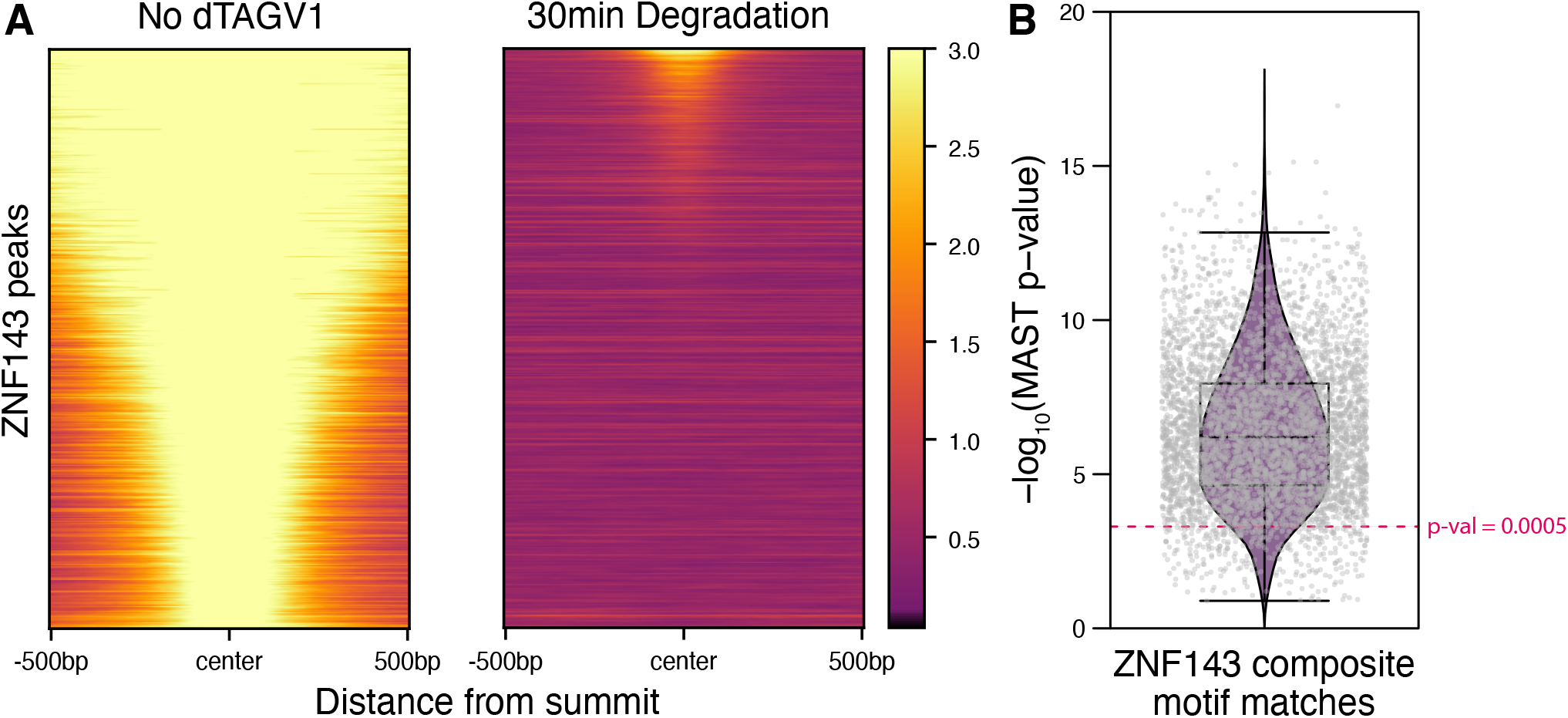
ZNF143 is rapidly depleted from chromatin after 30 minutes of dTAG treatment and we precisely define the ZNF143 binding site with motif analysis. Motif analysis found that all ZNF143 ChIP-seq peaks have a ZNF143 sequence that conforms to the Figure 1D composite motif with a p-value of 0.13 or less, with 93% of peaks at a p-value less than 0.0005.

**Fig. S2.**
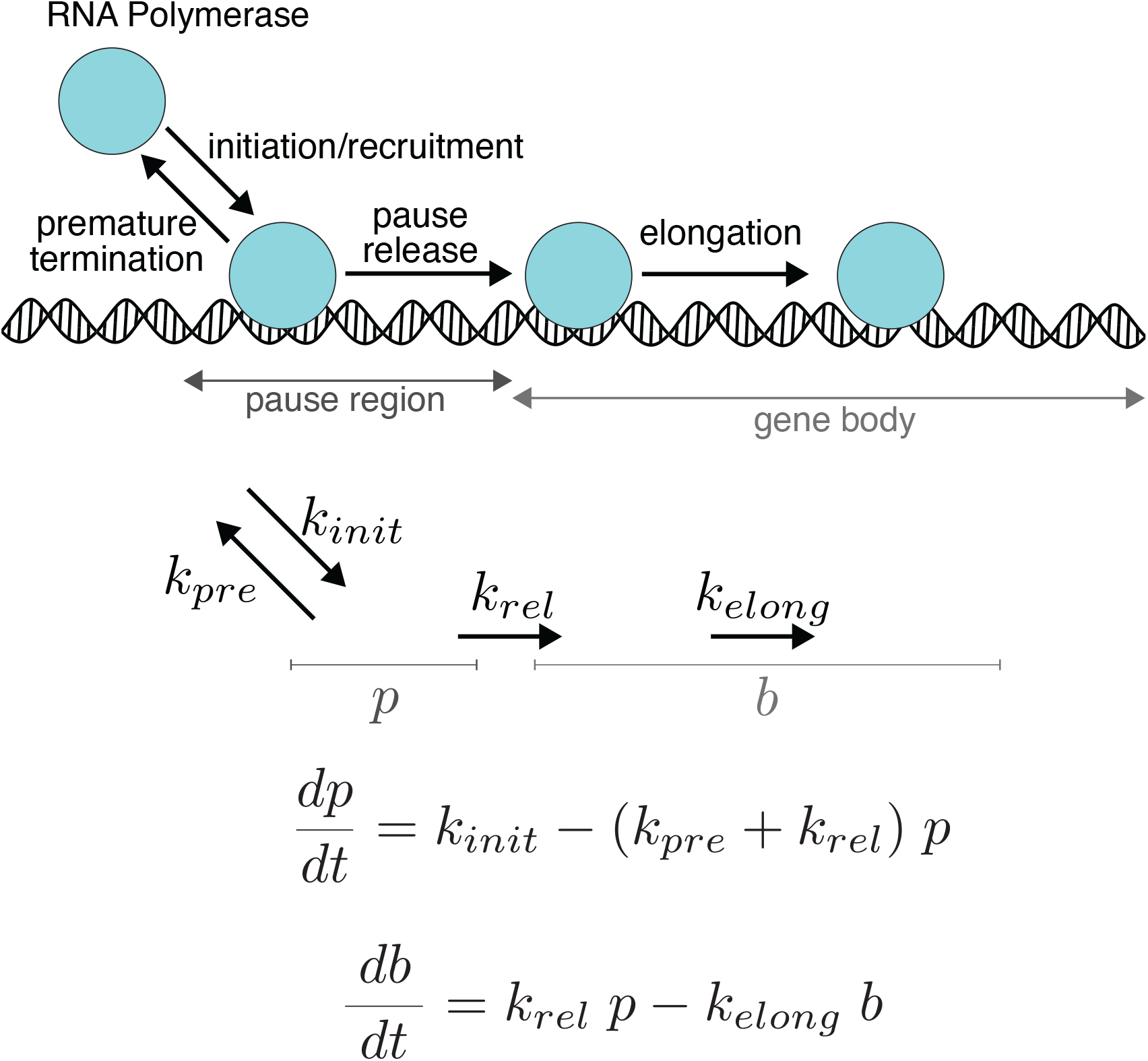
Compartment model for PRO-seq densities. This model and illustration was adapted from our previous work (Dutta et al. 2023). The pause densities *p* and body densities *b* are calculated directly from normalized PRO-seq data and elongation rates are constrained to a rate of 2-4kb/minute. The other parameters are calculated from the model.

**Fig. S3.**
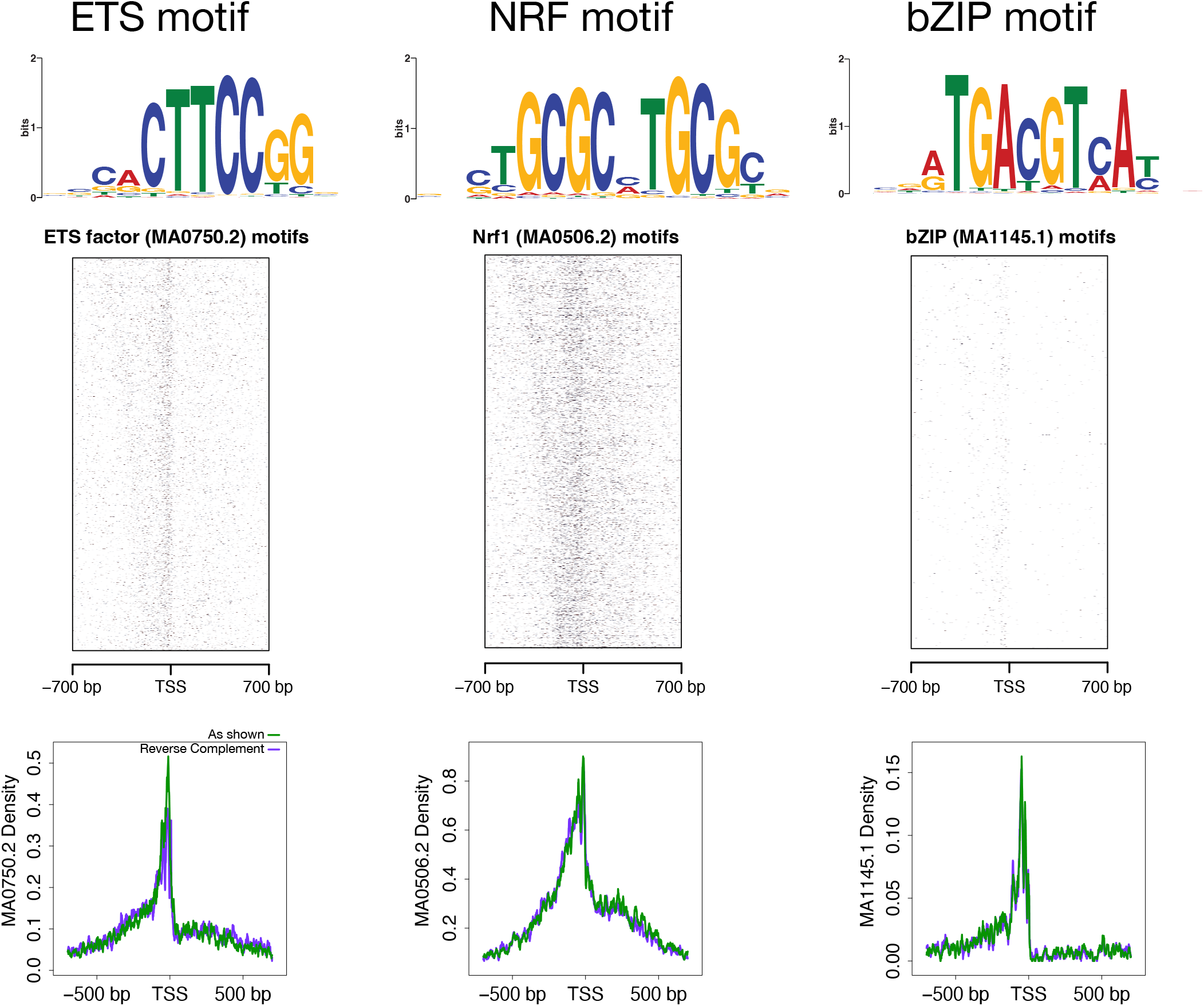
ETS, NRF, and bZIP motifs are enriched in promoters. *De novo* motif analysis within promoters revealed several motifs enriched between the same divergent and sense TSSs shown in Figure 7A. We used the best match to the JASPAR database (seqLogos) for each motif to identify their locations relative to TSSs with FIMO. The density plots quantify the distribution of the seqLogo-illustrated motif (top) and the reverse complement motif relative to TSSs.

**Fig. S4.**
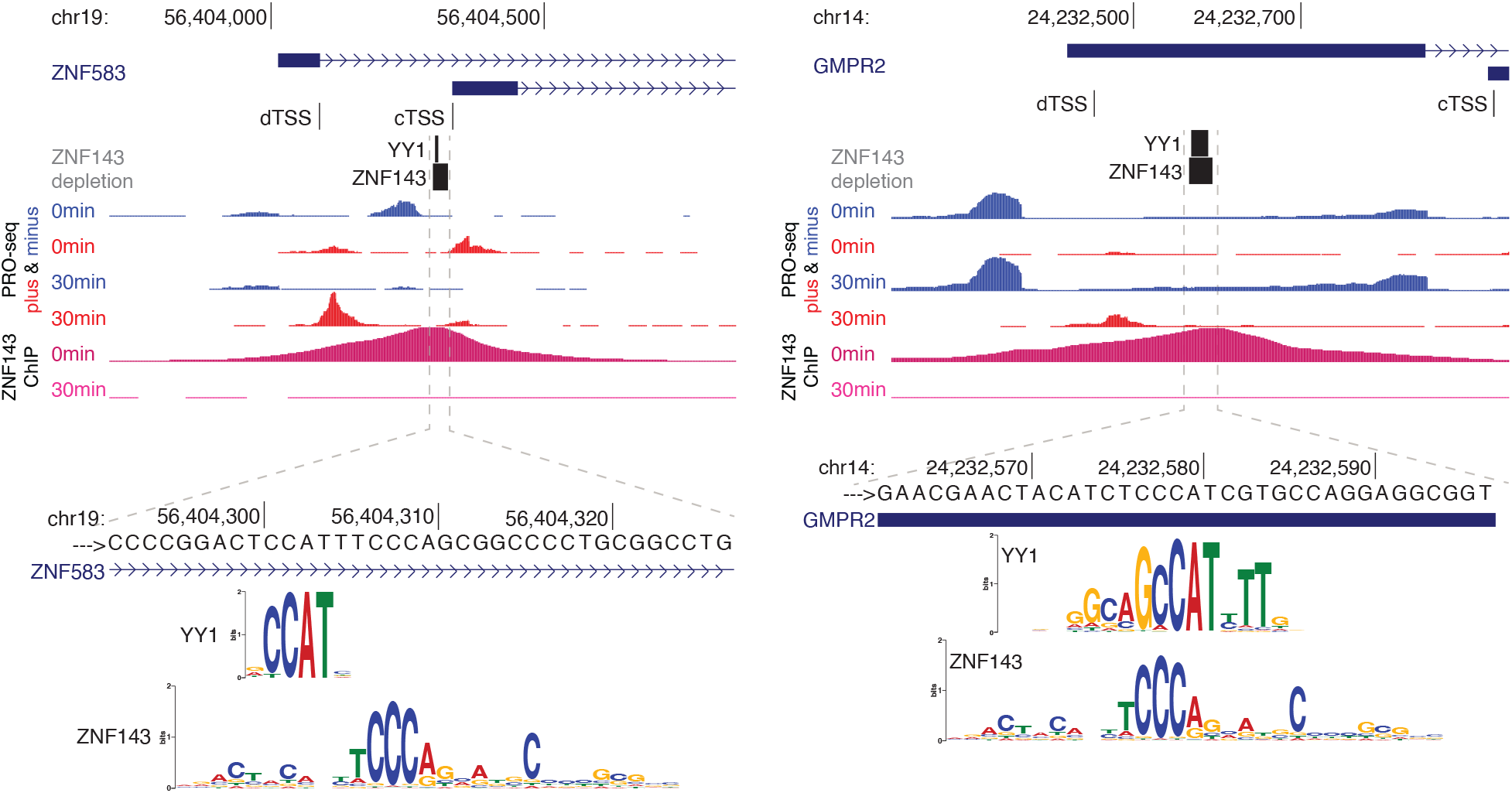
ZNF143 may block YY1 to reduce gene expression. The max TSS position of *ZNF583* and *GMPR2* shift upstream, suggesting a mechanism whereby YY1 access after ZNF143 degradation facilitates initiation at the dTAG TSS (cTSS: control max TSS position. dTSS: dTAG max TSS position).

**Fig. S5.**
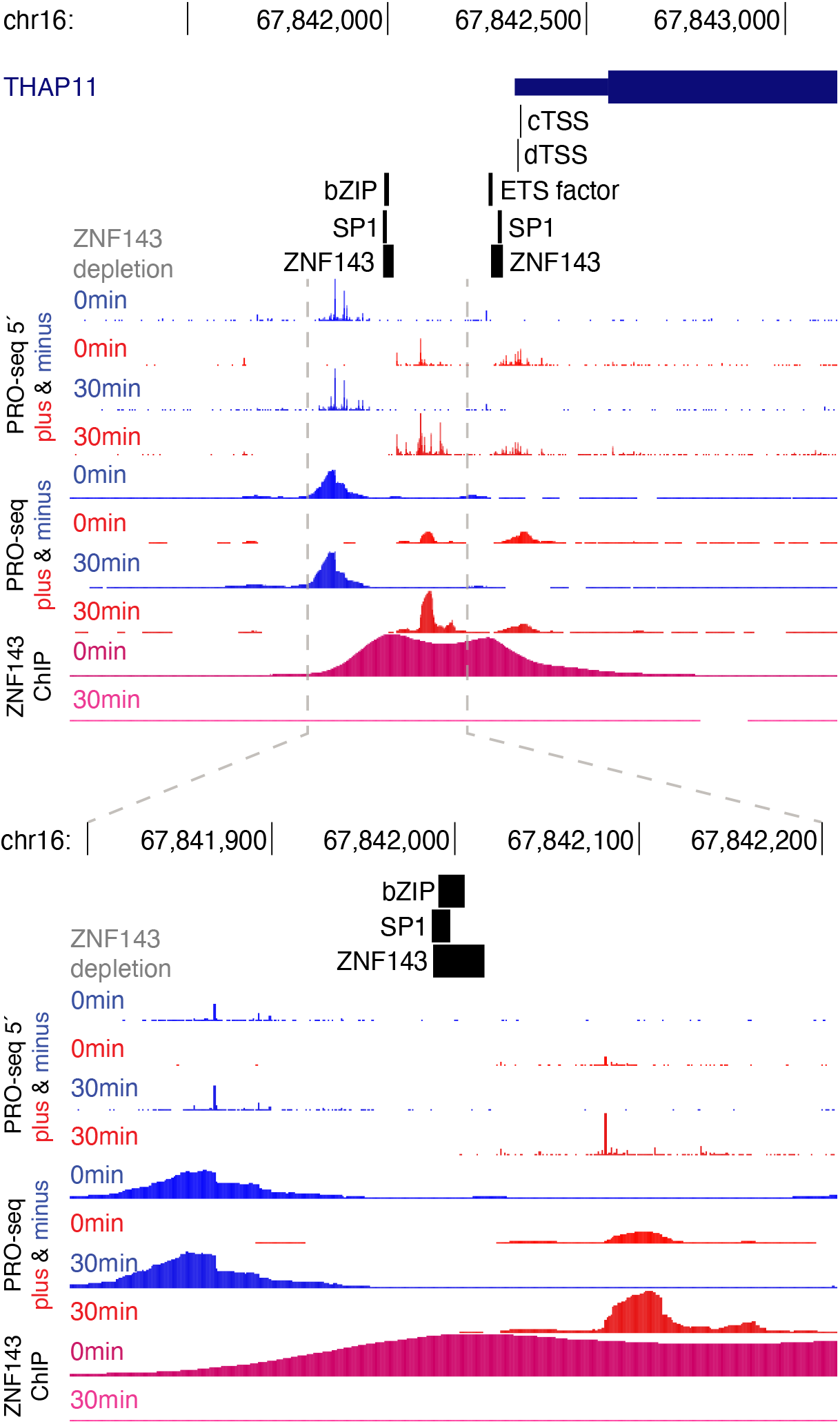
ZNF143 regulates *Thap11* via a complex mechanism. Depletion of ZNF143 allows for greater usage of a TSS upstream of both the control and dTAG TSSs. ZNF143 may repress transcription of *Thap11* by displacing other factors, such as ETS or a bZIP factor, that may be more potent activators of *Thap11*. Note that the ZNF143 motif shown in Figure 7D is most proximal to the TSS, but an upstream ZNF143 binding site/motif overlaps a bZIP motif. We do not conclude that SP1 redistributes because we did not observe a notable change in SP1 ChIP peak intensity.

## Bibliography

Ali MZ, Guharajan S, Parisutham V, and Brewster RC. 2023. Regulatory properties of transcription factors with diverse mechanistic function. bioRxiv pp. 2023–06.

Anderson WD, Duarte FM, Civelek M, and Guertin MJ. 2020. Defining data-driven primary transcript annotations with primarytranscriptannotation in r. Bioinformatics 36: 2926–2928.

Athanikar JN, Badge RM, and Moran JV. 2004. A yy1-binding site is required for accurate human line-1 transcription initiation. Nucleic acids research 32: 3846–3855.

Azofeifa JG and Dowell RD. 2017. A generative model for the behavior of rna polymerase. Bioinformatics 33: 227–234.

Bailey SD, Zhang X, Desai K, Aid M, Corradin O, Cowper-Sallari R, Akhtar-Zaidi B, Scacheri PC, Haibe-Kains B, and Lupien M. 2015a. Znf143 provides sequence specificity to secure chromatin interactions at gene promoters. Nature communications 6: 6186.

Bailey TL. 2021. STREME: accurate and versatile sequence motif discovery. Bioinformatics 37: 2834–2840.

Bailey TL and Gribskov M. 1998. Combining evidence using p-values: application to sequence homology searches. Bioinformatics 14: 48–54.

Bailey TL, Johnson J, Grant CE, and Noble WS. 2015b. The meme suite. Nucleic acids research 43: W39–W49.

Bates LE, Alves MR, and Silva JC. 2021. Auxin-degron system identifies immediate mechanisms of oct4. Stem Cell Reports 16: 1818–1831.

Benner C, Konovalov S, Mackintosh C, Hutt KR, Stunnenberg R, and Garcia-Bassets I. 2013. Decoding a signature-based model of transcription cofactor recruitment dictated by cardinal cis-regulatory elements in proximal promoter regions. PLoS genetics 9: e1003906.

Burke TW and Kadonaga JT. 1997. The downstream core promoter element, dpe, is conserved fromdrosophila to humans and is recognized by tafii60 of drosophila. Genes & development 11: 3020–3031.

Chen H, Lin RJ, Schiltz RL, Chakravarti D, Nash A, Nagy L, Privalsky ML, Nakatani Y, and Evans RM. 1997. Nuclear receptor coactivator actr is a novel histone acetyltransferase and forms a multimeric activation complex with p/caf and cbp/p300. Cell 90: 569–580.

Core LJ, Martins AL, Danko CG, Waters CT, Siepel A, and Lis JT. 2014. Analysis of nascent rna identifies a unified architecture of initiation regions at mammalian promoters and enhancers. Nature genetics 46: 1311–1320.

Core LJ, Waterfall JJ, and Lis JT. 2008. Nascent rna sequencing reveals widespread pausing and divergent initiation at human promoters. Science 322: 1845–1848.

DelRosso N, Tycko J, Suzuki P, Andrews C, Aradhana Mukund A, Liongson I, Ludwig C, Spees K, Fordyce P, et al. 2023. Large-scale mapping and mutagenesis of human transcriptional effector domains. Nature 616: 365–372.

Dong J and Guertin MJ. 2024. Infer transcription start sites from 5 prime ends of pro-seq reads. https://github.com/guertinlab/TSSinference.

Duarte FM, Fuda NJ, Mahat DB, Core LJ, Guertin MJ, and Lis JT. 2016. Transcription factors gaf and hsf act at distinct regulatory steps to modulate stress-induced gene activation. Genes & development 30: 1731–1746.

Dudnyk K, Cai D, Shi C, Xu J, and Zhou J. 2024. Sequence basis of transcription initiation in the human genome. Science 384: eadj0116.

Dutta AB, Lank DS, Przanowska RK, Przanowski P, Wang L, Nguyen B, Walavalkar NM, Duarte FM, and Guertin MJ. 2023. Kinetic networks identify twist2 as a key regulatory node in adipogenesis. Genome Research 33: 314–331.

Fondell JD, Ge H, and Roeder RG. 1996. Ligand induction of a transcriptionally active thyroid hormone receptor coactivator complex. Proceedings of the National Academy of Sciences 93: 8329–8333.

Fuda NJ, Ardehali MBf, and Lis JT. 2009. Defining mechanisms that regulate rna polymerase ii transcription in vivo. Nature 461: 186–192.

Gambi G, Di Simone E, Basso V, Ricci L, Wang R, Verma A, Elemento O, Ponzoni M, Inghirami G, Icardi L, et al. 2019. The transcriptional regulator sin3a contributes to the oncogenic potential of stat3. Cancer research 79: 3076–3087.

Gill G, Pascal E, Tseng ZH, and Tjian R. 1994. A glutamine-rich hydrophobic patch in transcription factor sp1 contacts the dtafii110 component of the drosophila tfiid complex and mediates transcriptional activation. Proceedings of the National Academy of Sciences 91: 192–196.

Gordon A. 2010. Fastx toolkit. https://github.com/agordon/fastx_toolkit.

Grandi FC, Modi H, Kampman L, and Corces MR. 2022. Chromatin accessibility profiling by atac-seq. Nature protocols 17: 1518–1552.

Grant CE, Bailey TL, and Noble WS. 2011. Fimo: scanning for occurrences of a given motif. Bioinformatics 27: 1017–1018.

Grøntved L, Waterfall JJ, Kim DW, Baek S, Sung MH, Zhao L, Park JW, Nielsen R, Walker RL, Zhu YJ, et al. 2015. Transcriptional activation by the thyroid hormone receptor through ligand-dependent receptor recruitment and chromatin remodelling. Nature communications 6: 7048.

Guertin M, Petesch S, Zobeck K, Min I, and Lis J. 2010. Drosophila heat shock system as a general model to investigate transcriptional regulation. In Cold Spring Harbor symposia on quantitative biology, volume 75, pp. 1–9. Cold Spring Harbor Laboratory Press.

Guertin MJ and Lis JT. 2010. Chromatin landscape dictates hsf binding to target dna elements. PLoS genetics 6: e1001114.

Guertin MJ, Zhang X, Coonrod SA, and Hager GL. 2014. Transient estrogen receptor binding and p300 redistribution support a squelching mechanism for estradiol-repressed genes. Molecular endocrinology 28: 1522–1533.

Hasterok S, Scott TG, Roller DG, Spencer A, Dutta AB, Sathyan KM, Frigo DE, Guertin MJ, and Gioeli D. 2023. The androgen receptor does not directly regulate the transcription of dna damage response genes. Molecular Cancer Research 21: 1329–1341.

Heidari N, Phanstiel DH, He C, Grubert F, Jahanbani F, Kasowski M, Zhang MQ, and Snyder MP. 2014. Genome-wide map of regulatory interactions in the human genome. Genome research 24: 1905–1917.

Hörlein AJ, Näär AM, Heinzel T, Torchia J, Gloss B, Kurokawa R, Ryan A, Kamei Y, Söderström M, Glass CK, et al. 1995. Ligand-independent repression by the thyroid hormone receptor mediated by a nuclear receptor co-repressor. Nature 377: 397–404.

Jindal GA, Bantle AT, Solvason JJ, Grudzien JL, D’Antonio-Chronowska A, Lim F, Le SH, Song BP, Ragsac MF, Klie A, et al. 2023. Single-nucleotide variants within heart enhancers increase binding affinity and disrupt heart development. Developmental Cell 58: 2206–2216.

Johnson AD, Meyer BJ, and Ptashne M. 1979. Interactions between dna-bound repressors govern regulation by the λ phage repressor. Proceedings of the National Academy of Sciences 76: 5061–5065.

Jones T, Sigauke RF, Sanford L, Taatjes DJ, Allen MA, and Dowell RD. 2024. A transcription factor (tf) inference method that broadly measures tf activity and identifies mechanistically distinct tf networks. bioRxiv pp. 2024–03.

Judd J, Wojenski LA, Wainman LM, Tippens ND, Rice EJ, Dziubek A, Villafano GJ, Wissink EM, Versluis P, Bagepalli L, et al. 2020. A rapid, sensitive, scalable method for precision run-on sequencing (pro-seq). Biorxiv pp. 2020–05.

Kent WJ, Sugnet CW, Furey TS, Roskin KM, Pringle TH, Zahler AM, and Haussler D. 2002. The human genome browser at ucsc. Genome research 12: 996–1006.

Kent WJ, Zweig AS, Barber G, Hinrichs AS, and Karolchik D. 2010. Bigwig and bigbed: enabling browsing of large distributed datasets. Bioinformatics 26: 2204–2207.

Kim Y, You HJ, Park SH, Kim MS, Chae H, Park J, Jekarl DW, Kim J, Kwon A, Choi H, et al. 2019. A mutation in znf143 as a novel candidate gene for endothelial corneal dystrophy. Journal of Clinical Medicine 8: 1174.

Kumar V and Chambon P. 1988. The estrogen receptor binds tightly to its responsive element as a ligand-induced homodimer. Cell 55: 145–156.

Kumar V, Green S, Stack G, Berry M, Jin JR, and Chambon P. 1987. Functional domains of the human estrogen receptor. Cell 51: 941–951.

Kwak H, Fuda NJ, Core LJ, and Lis JT. 2013. Precise maps of rna polymerase reveal how promoters direct initiation and pausing. Science 339: 950–953.

Langmead B and Salzberg SL. 2012. Fast gapped-read alignment with bowtie 2. Nature methods 9: 357–359.

Lewis EB. 1978. A gene complex controlling segmentation in drosophila. Nature 276: 565–570.

Li H, Handsaker B, Wysoker A, Fennell T, Ruan J, Homer N, Marth G, Abecasis G, and Durbin R. 2009. The sequence alignment/map format and samtools. Bioinformatics 25: 2078–2079.

Lim F, Solvason JJ, Ryan GE L. SH, Jindal GA, Steffen P, Jandu SK, and Farley EK. 2024. Affinity-optimizing enhancer variants disrupt development. Nature 626: 151–159.

Lin BC, Hong SH, Krig S, Yoh SM, and Privalsky ML. 1997. A conformational switch in nuclear hormone receptors is involved in coupling hormone binding to corepressor release. Molecular and cellular biology 17: 6131–6138.

Lin CY, Lovén J, Rahl PB, Paranal RM, Burge CB, Bradner JE, Lee TI, and Young RA. 2012. Transcriptional amplification in tumor cells with elevated c-myc. Cell 151: 56–67.

Liu S, Cao Y, Cui K, Tang Q, and Zhao K. 2022. Hi-trac reveals division of labor of transcription factors in organizing chromatin loops. Nature communications 13: 6679.

Lorenzin F, Benary U, Baluapuri A, Walz S, Jung LA, von Eyss B, Kisker C, Wolf J, Eilers M, and Wolf E. 2016. Different promoter affinities account for specificity in myc-dependent gene regulation. elife 5: e15161.

Love MI, Huber W, and Anders S. 2014. Moderated estimation of fold change and dispersion for rna-seq data with deseq2. Genome biology 15: 1–21.

Luse DS, Parida M, Spector BM, Nilson KA, and Price DH. 2020. A unified view of the sequence and functional organization of the human rna polymerase ii promoter. Nucleic acids research 48: 7767–7785.

Magnitov MD, Maresca M, Alonso Saiz N, Teunissen H, Braccioli L, and de Wit E. 2024. Znf143 is a transcriptional regulator of nuclear-encoded mitochondrial genes that acts independently of looping and ctcf. bioRxiv pp. 2024–03.

Mahat DB, Kwak H, Booth GT, Jonkers IH, Danko CG, Patel RK, Waters CT, Munson K, Core LJ, and Lis JT. 2016. Base-pair-resolution genome-wide mapping of active rna polymerases using precision nuclear run-on (pro-seq). Nature protocols 11: 1455–1476.

Martin M. 2011. Cutadapt removes adapter sequences from high-throughput sequencing reads. EMBnet. journal 17: 10–12.

Martins AL. 2014. R interface to query ucsc bigwig files. https://github.com/andrelmartins/bigWig.

Martins AL and Guertin MJ. 2018. Remove pcr duplicates from fastq files. https://github.com/guertinlab/fqdedup.

Martins AL, Walavalkar NM, Anderson WD, Zang C, and Guertin MJ. 2018. Universal correction of enzymatic sequence bias reveals molecular signatures of protein/dna interactions. Nucleic acids research 46: e9–e9.

McKnight SL and Kingsbury R. 1982. Transcriptional control signals of a eukaryotic proteincoding gene. Science 217: 316–324.

Meyer BJ, Maurer R, and Ptashne M. 1980. Gene regulation at the right operator (or) of bacteriophage λ: Ii. or1, or2, and or3: their roles in mediating the effects of repressor and cro. Journal of molecular biology 139: 163–194.

Muhar M, Ebert A, Neumann T, Umkehrer C, Jude J, Wieshofer C, Rescheneder P, Lipp JJ, Herzog VA, Reichholf B, et al. 2018. Slam-seq defines direct gene-regulatory functions of the brd4-myc axis. Science 360: 800–805.

Nabet B, Ferguson FM, Seong BKA, Kuljanin M, Leggett AL, Mohardt ML, Robichaud A, Conway AS, Buckley DL, Mancias JD, et al. 2020. Rapid and direct control of target protein levels with vhl-recruiting dtag molecules. Nature communications 11: 4687.

Nabet B, Roberts JM, Buckley DL, Paulk J, Dastjerdi S, Yang A, Leggett AL, Erb MA, Lawlor MA, Souza A, et al. 2018. The dtag system for immediate and target-specific protein degradation. Nature chemical biology 14: 431–441.

Narducci DN and Hansen AS. 2024. Putative looping factor znf143/zfp143 is an essential transcriptional regulator with no looping function. bioRxiv pp. 2024–03.

Ngondo-Mbongo RP, Myslinski E, Aster JC, and Carbon P. 2013. Modulation of gene expression via overlapping binding sites exerted by znf143, notch1 and thap11. Nucleic acids research 41: 4000–4014.

Nie Z, Hu G, Wei G, Cui K, Yamane A, Resch W, Wang R, Green DR, Tessarollo L, Casellas R, et al. 2012. c-myc is a universal amplifier of expressed genes in lymphocytes and embryonic stem cells. Cell 151: 68–79.

Nishimura K, Fukagawa T, Takisawa H, Kakimoto T, and Kanemaki M. 2009. An auxin-based degron system for the rapid depletion of proteins in nonplant cells. Nature methods 6: 917–922.

Nüsslein-Volhard C and Wieschaus E. 1980. Mutations affecting segment number and polarity in drosophila. Nature 287: 795–801.

Ptashne M. 2004. A genetic switch: phage lambda revisited .

Pupavac M, Watkins D, Petrella F, Fahiminiya S, Janer A, Cheung W, Gingras AC, Pastinen T, Muenzer J, Majewski J, et al. 2016. Inborn error of cobalamin metabolism associated with the intracellular accumulation of transcobalamin-bound cobalamin and mutations in znf143, which codes for a transcriptional activator. Human mutation 37: 976–982.

Reddy TE, Pauli F, Sprouse RO, Neff NF, Newberry KM, Garabedian MJ, and Myers RM. 2009. Genomic determination of the glucocorticoid response reveals unexpected mechanisms of gene regulation. Genome research 19: 2163–2171.

Sabo A, Kress TR, Pelizzola M, De Pretis S, Gorski MM, Tesi A, Morelli MJ, Bora P, Doni M, Verrecchia A, et al. 2014. Selective transcriptional regulation by myc in cellular growth control and lymphomagenesis. Nature 511: 488–492.

Santana JF, Collins GS, Parida M, Luse DS, and Price DH. 2022. Differential dependencies of human rna polymerase ii promoters on tbp, taf1, tfiib and xpb. Nucleic acids research 50: 9127–9148.

Sathyan KM, McKenna BD, Anderson WD, Duarte FM, Core L, and Guertin MJ. 2019. An improved auxin-inducible degron system preserves native protein levels and enables rapid and specific protein depletion. Genes & development 33: 1441–1455.

Sathyan KM, Scott TG, and Guertin MJ. 2020. Arf-aid: A rapidly inducible protein degradation system that preserves basal endogenous protein levels. Current protocols in molecular biology 132: e124.

Schmidt SF, Larsen BD, Loft A, and Mandrup S. 2016. Cofactor squelching: Artifact or fact? Bioessays 38: 618–626.

Schmidt SF, Larsen BD, Loft A, Nielsen R, Madsen JGS, and Mandrup S. 2015. Acute tnf-induced repression of cell identity genes is mediated by nfκb-directed redistribution of cofactors from super-enhancers. Genome research 25: 1281–1294.

Scholes C, DePace AH, and Sánchez Á. 2017. Combinatorial gene regulation through kinetic control of the transcription cycle. Cell systems 4: 97–108.

Schuster C, Myslinski E, Krol A, and Carbon P. 1995. Staf, a novel zinc finger protein that activates the rna polymerase iii promoter of the selenocysteine trna gene. The EMBO journal 14: 3777–3787.

Scott TG, Martins AL, and Guertin MJ. 2022. Processing and evaluating the quality of genome-wide nascent transcription profiling libraries. Biorxiv pp. 2022–12.

Scott TG, Sathyan KM, Gioeli D, and Guertin MJ. 2024. Trps1 modulates chromatin accessibility to regulate estrogen receptor alpha (er) binding and er target gene expression in luminal breast cancer cells. PLoS genetics 20: e1011159.

Seto E, Shi Y, and Shenk T. 1991. Yy1 is an initiator sequence-binding protein that directs and activates transcription in vitro. Nature 354: 241–245.

Sheppard HE, Dall’Agnese A, Park WD, Shamim MH, Dubrulle J, Johnson HL, Stossi F, Cogswell P, Sommer J, Levy J, et al. 2021. Targeted brachyury degradation disrupts a highly specific autoregulatory program controlling chordoma cell identity. Cell Reports Medicine 2.

Stengel KR, Ellis JD, Spielman CL, Bomber ML, and Hiebert SW. 2021. Definition of a small core transcriptional circuit regulated by aml1-eto. Molecular cell 81: 530–545.

Takahashi K and Yamanaka S. 2006. Induction of pluripotent stem cells from mouse embryonic and adult fibroblast cultures by defined factors. cell 126: 663–676.

Vinckevicius A, Parker JB, and Chakravarti D. 2015. Genomic determinants of thap11/znf143/hcfc1 complex recruitment to chromatin. Molecular and cellular biology 35: 4135–4146.

Wagner EJ, Tong L, and Adelman K. 2023. Integrator is a global promoter-proximal termination complex. Molecular cell 83: 416–427.

Walz S, Lorenzin F, Morton J, Wiese KE, von Eyss B, Herold S, Rycak L, Dumay-Odelot H, Karim S, Bartkuhn M, et al. 2014. Activation and repression by oncogenic myc shape tumour-specific gene expression profiles. Nature 511: 483–487.

Wang Z, Chu T, Choate LA, and Danko CG. 2019. Identification of regulatory elements from nascent transcription using dreg. Genome research 29: 293–303.

Westwood JT, Clos J, and Wu C. 1991. Stress-induced oligomerization and chromosomal relocalization of heat-shock factor. Nature 353: 822–827.

Yang Y, Zhang R, Singh S, and Ma J. 2017. Exploiting sequence-based features for predicting enhancer–promoter interactions. Bioinformatics 33: i252–i260.

Zhang K, Li N, Ainsworth RI, and Wang W. 2016. Systematic identification of protein combinations mediating chromatin looping. Nature communications 7: 12249.

Zhang M, Huang H, Li J, and Wu Q. 2024. Znf143 deletion alters enhancer/promoter looping and ctcf/cohesin geometry. Cell Reports 43.

Zhang Y, Liu T, Meyer CA, Eeckhoute J, Johnson DS, Bernstein BE, Nusbaum C, Myers RM, Brown M, Li W, et al. 2008. Model-based analysis of chip-seq (macs). Genome biology 9: 1–9.

